# A unified photosensitizer platform for *in situ* DNA-, RNA-, and protein-directed proximity labeling

**DOI:** 10.64898/2026.04.30.721698

**Authors:** Elijah B. Biletch, Conor P. Herlihy, Lidan Li, Erin Kim, Mary Krebs, Conor J. Kelly, Nicolas J. Longhi, Olivia Weissenfels, Hanna G. Goldberg, Kyle Brandt, Jonathan B. Grimm, Luke D. Lavis, Edward L. Huttlin, Melody M. H. Li, Devin K. Schweppe, Keriann M. Backus, Brian J. Beliveau

## Abstract

Cells depend on the spatial organization of proteins, RNA, and DNA into discrete subcellular compartments. Previous methods have largely centered on measuring spatial organization based on only one of these biomolecular classes at a time. Here, we demonstrate that POCA photocatalytic proximity labeling can serve as a unified photosensitizer-based platform for profiling the proximal proteomes of protein, RNA, and DNA targets within a single experimental framework. We show that POCA can harness standard immunofluorescence or *in situ* hybridization workflows to specifically target organic fluorophore photosensitizers to intracellular targets for proximity labeling in fixed cells. POCA-targeted proximity labeling requires minimal cellular input and does not require genetic engineering. Additionally, POCA photosensitizers are selected to also be fluorescent, enabling direct confirmation of on-target localization by imaging prior to proteomic analysis. To demonstrate broad utility, we apply POCA across multiple molecular targets spanning protein, RNA, and genomic DNA, including components of the nuclear pore complex, nucleolus, nuclear speckles, telomeres, and pericentromeric heterochromatin. By anchoring proximity labeling to both a protein and an RNA within the same nuclear compartment, we resolve shared and distinct proximal proteomes from orthogonal molecular perspectives.

## Introduction

Spatial organization of biomolecules is essential for cellular and organismal function. For example, metabolic proteins are partitioned within mitochondria; diverse cell signaling pathways coordinate at the plasma membrane; and chromatin is densely packed into transcriptionally active and silent compartments within the nucleus whose regulatory elements function across large genomic distances to contact the genes they control^1^. Beyond these classical examples, membraneless condensates, phase-separated assemblies, and transient microenvironments concentrate specific molecules in response to cellular signals. Phase-separated bodies extend the organizational logic of the cell to structures that lack a bounding membrane^2^. For example, the nucleolus concentrates machinery for ribosome biogenesis, and nuclear speckles recruit splicing factors involved in pre-mRNA maturation^2–8^. Characterizing subcellular structures has demonstrated the importance of their organization. For example, proteomic surveys of the nucleolus revealed unexpected connections to DNA damage response and cell cycle control^9,10^ and studies of stress granule composition identified FUS and TDP-43 as resident proteins^11,12^. Mislocalization and aggregation of FUS and TPD-43 have since been identified as hallmarks of ALS and frontotemporal dementia, drawing into question their functional relevance within stress granules^13,14^. Subcellular ‘omics’ approaches^15–17^ have already had considerable impact revealing regulatory principles that underlie both normal cellular function and their disruption in disease.

Proximity labeling is one such spatial method used to define subcellular molecular composition in an unbiased manner^18,19^. Canonically, proximity labeling is achieved by a catalyst, such as an engineered biotin ligase for BioID or peroxidase for APEX2, tethered to a protein of interest to generate short-lived reactive intermediates that covalently modify neighboring biomolecules within a defined radius, most commonly with biotin. The labeled molecules can then be recovered by affinity enrichment and identified by mass spectrometry or sequencing. BioID^20^, based on a promiscuous bacterial biotin ligase, revealed unexpected components of the centrosome and centriolar satellites, several of which were subsequently shown to be mutated in ciliopathies^21^. APEX^22^, an ascorbate peroxidase, and its improved derivative APEX2^23^, resolved the proteome of the mitochondrial intermembrane space, a compartment too narrow and biochemically intractable for conventional purification, identifying dozens of previously uncharacterized resident proteins^24^. TurboID^25^, an evolved biotin ligase with labeling kinetics fast enough for *in vivo* applications, extended proximity labeling across a range of organisms and tissues, establishing that the approach could operate in intact biological systems without toxic reagents^26^. While powerful, these platforms require genetic engineering to express catalyst-bait fusion proteins, limiting application to systems amenable to genetic engineering and targets that tolerate fusion without disruption to their biology.

The requirement for engineered expression can be circumvented by recruiting the labeling catalyst through an antibody rather than a genetic fusion. Methods such as EMARS^27^, SPPLAT^28^, and BAR^29^ demonstrated this principle by using antibody-conjugated horseradish peroxidase (HRP) to label the proximal proteomes surrounding endogenous proteins in fixed cells and tissues. BAR established that antibody-delivered HRP could recover high-confidence proximal proteomes for lamin A/C across diverse cell types, including primary samples inaccessible to genetic engineering^29^. Subsequent work extended this targeting logic to nucleic acid targets, enabling identification of RNA- or chromatin-proximal proteins, DNA, and RNA, including our work developing DNA O-MAP and RNA O-MAP which use oligonucleotide-directed, *in situ* hybridization to recruit HRP to specific DNA loci and RNAs—e.g., *Xist* and the X-inactivation centers of the X chromosome^30–32^. Yap and colleagues established HyPro, an analogous approach using APEX2 to label proteins proximal to noncoding RNAs *45S*, *NEAT1*, and *PNCTR*^33^. These antibody- and hybridization-based proximity labeling methods map the proteomes of nuclear bodies and chromatin domains and demonstrated that standard immunofluorescence (IF) and *in situ* hybridization (ISH) workflows could effectively deliver proximity labeling catalysts. However, these methods are constrained by their use of protein catalysts which require special handling to maintain activity, exhibit batch-to-batch variation, have slower protein diffusion in crowded intracellular environments, and can be difficult to quality control^34,35^.

Photocatalytic proximity labeling has emerged as a complement to enzyme-based approaches, offering temporal control through light-activated chemistry and reactive intermediates with shorter half-lives than enzymatic systems^36–38^. The shorter half-lives in particular have the potential to reduce off-target proximal labeling by limiting diffusion of reactive intermediates. Platforms such as µMap^36^, MultiMap^37,39^, and LITag^40^ demonstrated the growing impact of this approach at the cell surface, using photocatalysts conjugated to antibodies or small molecules to map proximal regulators of cellular receptors, cell-cell synapses, and biotherapeutic target environments on live cells. µMap uses organometallic iridium complexes that generate reactive carbene intermediates through Dexter energy transfer, with half-lives on the order of nanoseconds^36^. This chemistry has since been extended to formalin fixed, paraffin embedded tissue sections, where antibody-conjugated µMap photocatalysts labeled cell surface CD20 and its proximal proteome on fixed lymphoma specimens^41^. Genetically encoded, singlet-oxygen-generating photosensitizers such as miniSOG^42^ extend the approach intracellularly by generating singlet oxygen upon light irradiation, which oxidizes proximal biomolecules—primarily histidine in proteins—to create reactive intermediates that can be biotinylated and enriched. However, similar to enzyme-based platforms, these approaches require genetic manipulation to deliver the catalyst which can be time consuming and expensive.

Fluorophore photosensitizers represent a mechanistically distinct and practically attractive class of PPL reagents^37,38,43^. These small organic molecules, including the Eosin Y photosensitizers used in MultiMap^39^, exploit a fundamental photophysical tradeoff between fluorescence for imaging and singlet oxygen generation for proximity labeling. Halogen substitution or related heavy atom effects increase the effect of intersystem crossing and singlet oxygen generation at the modest expense of fluorescence quantum yield. This tradeoff results in a molecule bright enough for imaging and efficient enough for singlet oxygen generation to drive proximity labeling chemistry^43^. Because the same molecule serves two functions, successful photosensitizer localization to the protein of interest can be confirmed by imaging before the sample is committed to mass spectrometry or sequencing. Photosensitizers can be readily conjugated to antibodies, oligonucleotides, and small molecules, enabling their delivery without the need for genetic engineering^37,38,44–46^. Despite this versatility, organic fluorophore photosensitizers have only been demonstrated in live cells under native conditions, and their compatibility with the fixation and permeabilization conditions required for ISH and IF targeting of intracellular DNA, RNA, and protein has not, to our knowledge, been established.

Building on our experience with HRP-conjugated probes, we investigated whether a small-molecule photosensitizer could replace the protein catalyst in a universal targeting platform. Such a platform would use organic fluorophores as photosensitizers, enable conjugation to diverse targeting molecules, and provide an imaging readout to confirm catalyst delivery, all without the need for genetic engineering. We recently described POCA (<u>p</u>hotosensitizer-dependent <u>o</u>xidation and <u>c</u>apture by <u>a</u>mine)^38^, a singlet oxygen-based photocatalytic approach that established dibromofluorescein (DBF) and the rhodamine analogue JF_570_ as effective intracellular photosensitizers and defined reaction conditions compatible with affinity purification and quantitative mass spectrometry. Our initial implementation used HaloTag fusions or direct lipid conjugates to recruit the photosensitizer and was designed to work in live cells. Here, we extend the POCA platform to endogenous DNA, RNA, and protein targets using photosensitizers conjugated to secondary antibodies and oligonucleotide probes deployed through standard IF and ISH workflows. We apply this approach to ten molecular targets including proteins, RNAs, and genomic loci. These targets span major compartments of the nucleus and cytoplasm, recovering proximal proteomes consistent with the known biology and new putative factors. Importantly, we can recover these proteomes starting from just two million cells per replicate. Cross-modality pairing of IF-POCA and ISH-POCA (protein and RNA targeting) enables direct local proteome comparisons between different biomolecule targets that occupy the same subcellular compartment, a capability we demonstrate in nucleoli and nuclear speckles. Thus, the POCA platform is a flexible, cost-effective, generalized framework for the spatial analysis of subcompartment composition.

## Results

### Establishing POCA photosensitizer-based proximity labeling in fixed cells

The ISH- and IF-POCA platform consists of a targeting module for specific target recognition and a labeling module for photosensitizer-based proximity labeling. The central design choice we chose to develop here was the use of secondary reagents as photosensitizer delivery vehicles for labeling (**Figure 1A**). In ISH-POCA, a photosensitizer-conjugated secondary oligonucleotide hybridizes to a primary oligonucleotide targeting an RNA or genomic DNA locus. In IF-POCA, a photosensitizer-conjugated secondary antibody binds a primary antibody targeting a protein of interest. Upon irradiation, the photosensitizer generates singlet oxygen that oxidizes nearby proteins, and a biotin-amine capture reagent covalently tags the oxidized residues, enabling enrichment and identification by mass spectrometry. Because the photosensitizer is carried on secondary reagents, a single conjugated secondary antibody or oligonucleotide can deliver POCA to any target for which a validated primary reagent exists. This means a small number of characterized secondary reagents serve as universal delivery vehicles, and each reagent only needs to be validated once to enable reuse across multiple experiments. Within this framework, an important prerequisite was to verify that the photosensitizer behaves as an effective fluorophore under ISH/IF conditions, maintaining sufficient brightness, stability, and signal-to-noise performance. DBF was selected for its combination of efficient singlet oxygen generation, sufficient fluorescence for imaging, and proven performance in proximity labeling workflows^47–51^. We conjugated DBF to secondary antibodies and oligonucleotide probes via NHS ester chemistry (**Figure S1A,B**) and validated its fluorophore properties in both ISH and IF applications (**Figure S1C,D**).

**Figure 1.**
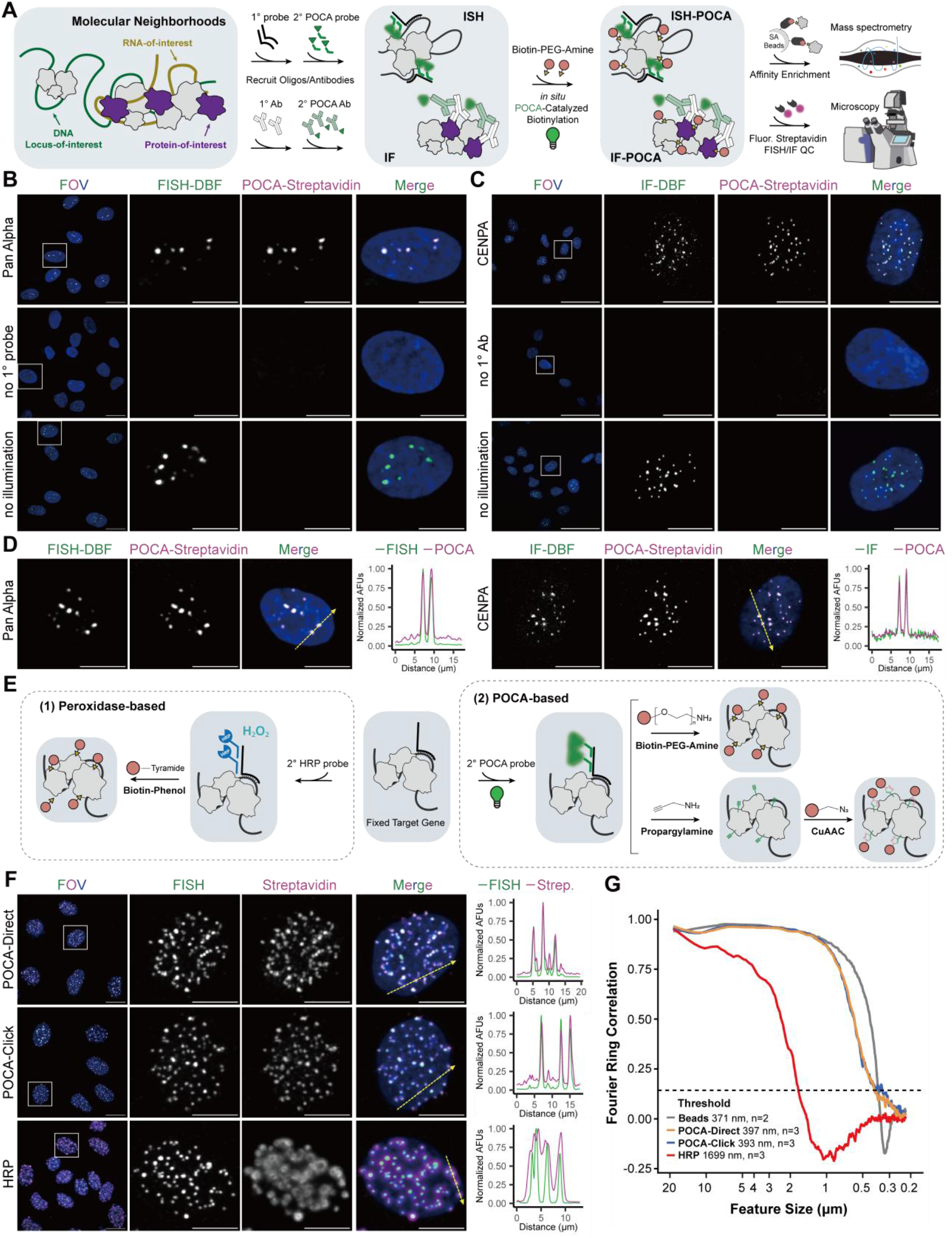
ISH- and IF-POCA labels nuclear structures. **A)** Schematic showing the workflow for ISH- and IF-POCA labeling experiments. **B)** Representative images of the pericentromeric Pan Alpha ISH-POCA labeling depicting FISH-DBF (green), POCA-Streptavidin (magenta) signal, a merged image of both labeling channels with DAPI (blue), and a cropped image of a single cell from the merged image. Overlap between DBF and streptavidin signal appears white. **C)** Representative images of CENPA IF-POCA labeling depicting IF-DBF (green), POCA-Streptavidin (magenta) signal, a merged image of both labeling channels with DAPI (blue), and a cropped image of a single cell from the merged image. Overlap between DBF and streptavidin signal appears white. **D)** Representative images as in **B** & **C**, with depiction of the line scan through two Pan Alpha or CENPA foci. Line scan graphs show fluorescence intensity of DBF (green) and streptavidin (magenta) along the dotted yellow lines in the adjacent representative images. **E)** Schematic depicting the HRP, direct POCA, and click POCA labeling approaches. **F)** Representative images of the minor satellite ISH-POCA labeling, as in **B** & **C**, in EY.T4 mouse embryonic fibroblast cells. Line scans show fluorescence intensity of FISH (green) and streptavidin (magenta) along the dotted yellow lines for each of the three labeling approaches. **G)** Mean Fourier Ring Correlation (FRC) across a range of feature sizes for the three labeling approaches compared to TetraSpeck 200 nm fluorescent beads (Beads) as a measure of the microscope’s optical limit. Each line is the mean FRC across three fields of view (each containing 6–21 nuclei and 381–1273 puncta) for each labeling approach targeting minor satellites in EY.T4 mouse embryonic fibroblasts cells, as shown in **F**. Dotted line represents the 1/7 FRC threshold^54^. FRC analysis was performed on maximum intensity in Z projections.

To develop the POCA platform with DBF, we needed to identify an optimal set of conditions that enable correct target binding, stable fluorophore performance, efficient labeling, low background, and a simple operational workflow. For initial benchmarking, we used an ISH probe that targets the pericentromeric alpha satellite repeat family (‘Pan Alpha’), for which we have previously reported extensive imaging and HRP-based proximity labeling data^52^. We simultaneously monitored the ISH signal from DBF and biotin deposition via fluorescent streptavidin labeling using fluorescence microscopy to define effective ISH-POCA labeling conditions. We then performed a systematic evaluation of key parameters, including biotin-amine concentration, light intensity, and irradiation time using commercially available light sources (**Table S1**). Parameter optimization revealed 2 mM of biotin-amine to have a higher signal-to-noise ratio than lower concentrations (0.5-1 mM) with less off-target labeling signal than at a higher concentration (5 mM) (**Figure S2A**). For light intensity, a power density of ∼0.73 mW/cm² generated the best signal-to-noise ratio, with lower powers generating diminished labeling intensity (**Figure S2B**). We also investigated how the duration of the labeling reaction affected the biotin deposition signal. Labeling for 5 minutes resulted in excellent on-target intensity with the lowest off-target signal compared to 10 and 15 minute intervals in relation to the baseline FISH fluorescence (**Figure S2C**). In summary, optimal labeling was achieved using 2 mM biotin-amine with a power density of ∼0.73 mW/cm² for 5 minutes for ISH-POCA. The Pan Alpha fluorescence pattern obtained here matches the expected pattern seen in our prior imaging results^52^ (**Figure 1B**), and the immunofluorescence signal of CENPA exhibited a characteristic centromeric localization pattern (**Figure 1C**), further validating the specificity and robustness of the system.

A benefit of using fluorescent photocatalysts for proximity labeling is the potential to perform target imaging and proximity labeling with the same molecule. To this effect, we confirmed that DBF retained fluorescence after the POCA irradiation step, enabling both photocatalytic labeling and direct imaging of the target in a single round of probe delivery (**Figure 1B**). Line-scan analysis across individual foci confirmed strong agreement between the target signal and biotin deposition (**Figure 1D**). This dual functionality allows co-visualization of the target and fluorescent streptavidin-detected biotinylation *in situ*, providing quality control of labeling specificity before samples are committed to downstream proteomic analysis., e.g., mass spectrometry.

To establish the broad applicability of ISH- and IF-POCA, we validated labeling with a panel of DNA, RNA, and protein targets across cell context and species. For DNA, we labeled pericentromeric alpha satellites in both hTERT-RPE1 adherent and K562 suspension cells to extend applicability beyond adherent cultures; minor satellite repeats in EY.T4 mouse embryonic fibroblasts to confirm cross-species utility; and the single-copy ∼83 kb *HOXD* locus in hTERT-RPE1 cells to test single-locus labeling sensitivity (**Figure S3A**). ISH-POCA labeling across a diverse set of DNA (Pan Alpha, minor satellites, *HOXD*) and RNA (*XIST*, *CBX5*, *MALAT1*, and *ITS1*) targets mirrored FISH signal regardless of target size, cell type, or species (**Figure S3A-C**). Additionally, IF-POCA labeling using DBF-conjugated secondary antibodies against either rabbit or mouse primaries resolved CENPA at centromeres, TOM20 at the mitochondrial outer membrane, and COXIV at the mitochondrial inner membrane, with the latter two demonstrating applicability beyond the nucleus to cytoplasmic targets (**Figure S3D**). For both IF-POCA targeting of CENPA and ISH-POCA targeting of Pan Alpha, omitting either the primary reagent or illumination eliminated biotin deposition (**Figure 1B,C**).

In light of the optimized labeling conditions and generalizable application of POCA labeling for proteins or nucleic acids with biotin-amine, we wanted to confirm the efficiency of direct biotin deposition chemistries compared to our previous usage of a click-based biotinylation after initial POCA labeling with propargylamine^38^. Because POCA operates in fixed, permeabilized cells, both cell-impermeable biotin-amine and cell-permeable propargylamine (conjugated to biotin-azide via click chemistry) can serve as the capture reagents (**Figure 1E**). We compared both POCA biotinylation strategies to canonical peroxidase-based biotin deposition using an ISH probe targeting centromeric mouse minor satellite repeats in EY.T4 mouse embryonic fibroblast cells, which we have previously benchmarked^52,53^. Line scan analysis across individual foci showed that biotin signal tracks DBF fluorescence closely and drops off sharply outside the target for both direct and click-mediated deposition by POCA (**Figure 1F**). POCA-based labeling generated narrower biotin deposition more consistent with minor satellite FISH when compared to HRP-based labeling (**Figure 1F**).

We next asked whether POCA biotinylation faithfully recapitulates the spatial distribution of the FISH probe signal down to the optical resolution limit. Fourier Ring Correlation (FRC) is widely used in cryo-electron microscopy and super-resolution imaging to estimate the resolution of a reconstruction by splitting a dataset into two independent halves and measuring their agreement at each spatial frequency^54,55^. All labeling approaches agree at coarse spatial scales. The distinguishing feature is how fine a spatial scale that agreement extends to before it breaks down. We reasoned that the same framework could be applied to evaluate proximity labeling by treating the FISH and streptavidin channels as two independent measurements of the same underlying spatial structure. FRC decomposes the two images into spatial scales—those encompassing large features such as whole nuclei down to fine features such as individual foci— and evaluates their agreement at each scale in a single operation rather than requiring repeated measurements at each length scale independently. We adopted the 1/7 threshold, a well-validated criterion from cryo-EM for distinguishing reproducible structural detail from noise^54^, to define the finest spatial scale at which biotin deposition still tracks the FISH probe signal. As a reference for the optical ceiling, we first computed FRC between the 488 nm and 640 nm channels on 200 nm fluorescent bead slides imaged with identical acquisition settings, establishing a crossing the 1/7 threshold at 371 nm (**Figure 1G**). Both POCA labeling approaches were indistinguishable from the bead optical limit above 1 µm and diverged only modestly at finer scales, crossing the 1/7 threshold at 397 ± 10 nm (direct biotinylation) and 393 ± 38 nm (click biotinylation)—within 26 nm of the 371 nm bead floor. In contrast, HRP-based biotinylation crossed the 1/7 threshold at 1699 ± 44 nm, a 4.3-fold larger length scale. Below approximately 1 µm, HRP FRC values became negative, indicating a biotin labeling pattern that is anticorrelated with the FISH probe signal at fine spatial scales, likely due to diffusion of tyramide radicals away from on-target HRP molecules. These data demonstrate that POCA achieves near-diffraction-limited spatial fidelity in proximity labeling, consistent with previous reports of improved spatial restriction with photocatalytic platforms^36,37^ relative to peroxidase-based approaches.

### DNA-targeted ISH-POCA recovers locus-specific proximal proteomes

Having established that POCA produces proximity-confined biotinylation across protein, RNA, and DNA targets, we next asked whether this labeling could be coupled to quantitative mass spectrometry for unbiased discovery of proximal proteomes. Our initial implementation of POCA for imaging used a set up that allowed us to photoilluminate a single 18-well chamber slide (40,000–50,000 cells per well). In order to apply POCA at the scale required for quantitative proteomics across multiple conditions, we sought to increase the amount of material labelled per experiment.

To overcome sample limitations, we constructed simple, modular photoillumination platforms in which light sources were positioned in a manner sufficient to illuminate each well of a 6-well plate or 60-mm dish, enabling simultaneous POCA labeling across all cells regardless of format (∼1 million cells per well in a 6-well dish or ∼3 million cells per 60-mm dish, **Figure S4A,B**). These setups could be readily assembled using commercially available components and adapted to different laboratory environments, as demonstrated by the two independent implementations of the platform used in this work (**Table S1**). Even at these larger scales, the cost for reagents used to biotinylate proteins was less than $3 per replicate (**Table S1**).

Using the new photoillumination platforms, we verified that DBF for proteomics performed as expected in our hands based on a strong preference for the biotinylation of histidines^38^ (**Figure S4C-F**). We then asked whether ISH-POCA could resolve the proximal proteomes of two well-characterized repetitive DNA loci, pericentromeric Pan Alpha and telomere repeat sequences in hTERT-RPE1 cells. We selected these regions given our success with targeting Pan Alpha with the POCA platform (**Figure 1B,D, Figure S3A**) and our previous success targeting both of these loci for oligo-based proximity labeling^32,52^. For each target we analyzed four replicates (each two wells of a 6-well plate) using ISH-POCA labeling and an additional well for imaging quality control to check the FISH signal and POCA labeling for each target (**Figure 2A,B**). We confirmed successful oligo labeling and biotin deposition based on FISH of DBF-oligos and fluorescent streptavidin staining of POCA-based biotinylation (**Figure 2C**); both were consistent with staining patterns previously seen for Pan Alpha and telomere FISH. Together these results demonstrate that we can achieve robust biotinylation with POCA in a large enough format (6-well plates) for quantitative proteomics.

**Figure 2.**
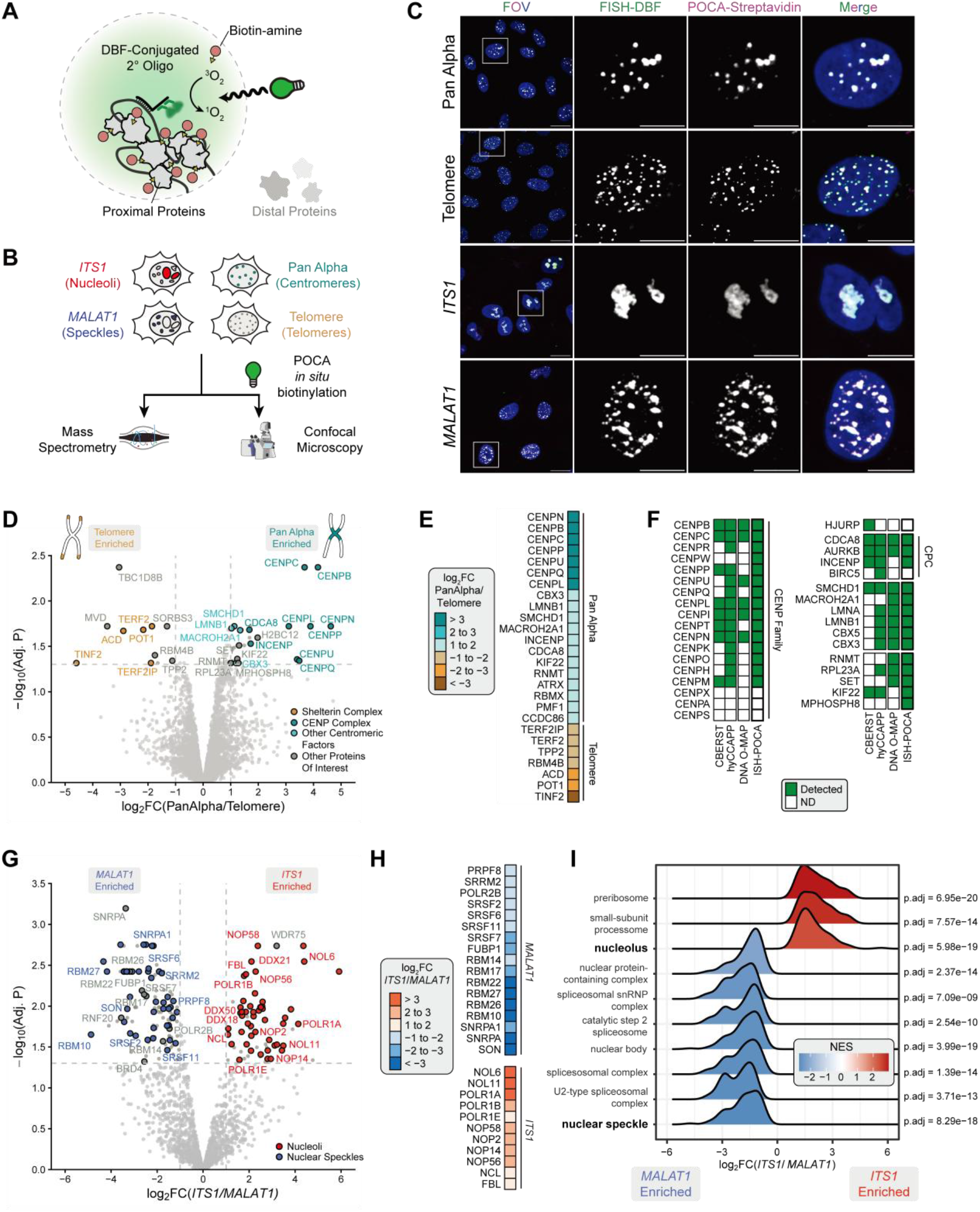
ISH-POCA reveals DNA and RNA proximal proteomes. **A)** Schematic showing the ISH-POCA labeling approach. **B)** Schematic showing the experimental workflow for ISH-POCA targeting distinct genomic loci and nuclear compartments. **C)** Representative images from each of the four ISH-POCA targets depicting FISH-DBF (green), POCA-Streptavidin (magenta), a merged image of both labeling channels plus DAPI (blue), and a cropped image of a single cell from the merged image. Overlap between DBF and Streptavidin signal appears as white. All images are maximum intensity projections. Scale bars are 20 µm in the standard images and 10 µm in the cropped images. **D)** Volcano plot of proteins identified at Pan Alpha satellites and telomeres. Each dot represents a single protein. Significantly enriched proteins associated with the shelterin and CENP complexes are labeled in tan and cyan, respectively. CENP and shelterin annotation lists were generated from the Human Protein Atlas (HPA)^15^, CORUM^80^, and UniProtKB^81^ databases. Other significantly enriched proteins of interest are denoted in grey. Hashed lines delineate the thresholds for significance, with greater than 2-fold change in relative enrichment and an FDR-corrected p-value less than 0.05. **E)** Heatmap summarizing log_2_ fold-changes for pairwise comparisons between Pan Alpha and telomere. **F)** Binary heatmap of proteins detected or not by Pan Alpha ISH-POCA and previous studies^32,100,101^. **G)** Volcano plot of proteins identified at *ITS1* and *MALAT1*. Each dot represents a single protein. Significantly enriched proteins associated with the nucleolus and nuclear speckles are labeled in red and blue, respectively. The nucleolar and nuclear speckle annotation lists were generated from the Human Protein Atlas (HPA)^15^, CORUM^80^, and UniProtKB^81^ databases. Hashed lines delineate the thresholds for significance, with greater than 2-fold change in relative enrichment and an FDR-corrected p-value less than 0.05. **H)** Heatmap summarizing log_2_ fold-changes for pairwise comparisons between *MALAT1* and *ITS1*. **I)** GSEA of significantly enriched GO terms from ISH-POCA of *MALAT1* and *ITS1* enriched proteomes. FDR-corrected p-values associated with each term are shown to the right. NES: normalized enrichment score. All raw and processed proteomics data associated with this figure can be found in **Table S5**.

We enriched biotinylated proteins from DNA-targeted ISH-POCA-labeled cells and characterized the resulting proximal proteomes by label-free, MS1-based quantitative proteomics (**Figure 2A,B**). DNA ISH-POCA proteomics identified 42 significantly enriched proteins comparing Pan Alpha and telomere proximal proteomes. Telomeres are uniquely bound by the six-subunit shelterin complex (TERF1, TERF2, TERF2IP, TINF2, ACD, POT1), which protects the single-stranded overhangs at the end of chromosomes^56–60^. Compared to Pan Alpha, telomeric ISH-POCA significantly enriched the shelterin complex, capturing five of the six shelterin complex members—TERF2, TERF2IP, TINF2, POT1, and ACD—an improvement over our previous work with DNA O-MAP^32^ (**Figure 2D,E, Figure S5A-C**). The recovery of intact protein complexes suggested that ISH-POCA preserves the interaction architecture of the labeled microenvironment. To assess this systematically, we asked whether the telomere and Pan Alpha ISH-POCA datasets were enriched for known protein-protein interactions beyond individual complex membership based on BioPlex^61,62^ interactome network (**Figure S5D**). Indeed, both datasets had highly significant levels of interactions between proteins enriched with ISH-POCA. Of these, telomere ISH-POCA enriched RNA binding and nuclear factors not previously annotated as telomeric, including RBM4B and TPP2 (**Figure 2E, S5B,E**). Both proteins localize to nuclei^15^, and, according to BioPlex, RBM4B interacts directly with the telomerase component DKC1^16^. Moreover, RBM4B protein abundance was significantly correlated with ACD/TPP1 and TERF2IP abundances in CCLE cell lines^63^. The convergence of these datasets strongly suggests that RBM4B and TPP2 are putative telomere proximal proteins.

Human centromeres are epigenetically defined genomic regions vital to proper chromosome segregation, kinetochore assembly, and spindle microtubule attachment^64–70^. The protein environment of mammalian centromeres is largely made up of the centromere protein (CENP) family members and the chromosome passenger complex^71–73^. As such, Pan Alpha ISH-POCA significantly enriched the CENP complex compared to telomere ISH-POCA (**Figure 2D,E, S5A,F**). Other centromere-associating factors were enriched at Pan Alpha, including chromosome passenger complex members INCENP and Borealin (CDCA8), and other centromere factors CBX3 (HP1*γ*), MacroH2A1, and SMCHD1. Known centromere localized proteins CBX5 (HP1ɑ, Log_2_(Pan Alpha/Telo)=0.881) and LMNA (Log_2_(Pan Alpha/Telo)=0.904) did not pass the stringent protein fold-change thresholds, but were statistically significant in Pan Alpha ISH-POCA. LMNA enrichment at Pan Alpha loci was expected due to known co-localization of pericentromeric heterochromatin and lamina-associated domains^74^. Pan Alpha also enriched proteins not previously annotated as centromeric, including RNMT, KIF22, MPHOSPH8, RPL23A and SET (**Figure 2E, S5G,H**). As with the telomeric analysis, proteins significant with Pan Alpha ISH-POCA were significantly enriched for protein interactions in BioPlex^16,61^, including those with known centromeric factors (**Figure S5G,H**). Corroborating the Pan Alpha ISH-POCA results, recent work demonstrated that knockout of MPHOSPH8 led to derepression of alpha satellites in human stem cells^75^. The enrichment of MPHOSPH8 with Pan Alpha ISH-POCA suggests that derepression could be through proximal, potentially direct, transcriptional silencing. These results demonstrate that ISH-POCA achieved locus-level resolution of chromatin proximal proteomes, recovering both canonical and putative factors to enable future investigation of these loci.

### RNA-targeted ISH-POCA recovers long non-coding RNA proximal proteomes

Based on the selectivity and sensitivity of DNA ISH-POCA, we wanted to test the application of ISH-POCA to measure proximal proteomes for spatially-distinct non-coding RNAs (ncRNAs) *ITS1* and *MALAT1*^30^. *ITS1* is a nucleolar-localized ncRNA from the transcribed spacer domain within the 47S pre-rRNA transcript^76,77^. *MALAT1* is a nuclear speckle-localized regulatory long ncRNA^78^ that acts as a transcriptional regulatory factor and molecular scaffold for ribonucleoprotein complexes^79^. Using ISH-POCA, we targeted *ITS1* and *MALAT1* in hTERT-RPE1 (2 million cells per replicate, **Figure 2A,B**). As with our DNA targets, we confirmed specificity of biotin deposition by microscopy based on the overlap of FISH and streptavidin staining (**Figure 2C**). From direct comparison of *ITS1* and *MALAT1* proximal proteomes, 70 proteins were significantly enriched at *ITS1* and 119 proteins were significantly enriched at *MALAT1* (**Figure 2G,H**). As with DNA ISH-POCA, these sets of proteins were strongly enriched for known protein-protein interactions and had highly significant levels of interactions between the enriched proteins (**Figure S5D**). Of the *ITS1* hits, 60 (85.7%) were annotated as nucleolar localized proteins in at least one of the following datasets: the Human Protein Atlas (HPA)^15^, CORUM^80^, and UniProtKB annotations^81^. For *MALAT1*, 78 protein hits (65.5%) were annotated as nuclear speckle localized, a proportion consistent with recent TurboID-based proximal proteomics of nuclear speckle proteins^82^.

*MALAT1* ISH-POCA enriched structural components of nuclear speckles including SON and SRRM2^83–85^, pre-mRNA splicing factor SRSF7 implicated in mRNA translation and transport^86–88^, and proteins involved in RNA splicing and export (SNRPA, SNRPA1, RBM22, PRPF8) (**Figure 2G,H**). *MALAT1* also enriched BRD4, the short isoform of which has been implicated in transcription and phase separation^89^. Additionally, BRD4 interacts with p-TEFb which is thought to function at nuclear speckles following CDK9 T-loop phosphorylation^90^, implicating BRD4 as a potential nuclear speckle protein. *MALAT1* ISH-POCA also enriched for the E3 ubiquitin ligase RNF20 which, similar to CDK9, is implicated in H2B ubiquitination. RNF20 and subsequent H2B ubiquitination have also been shown to play a role in transcription and mRNA splicing^91^, further suggesting RNF20 may have an unappreciated nuclear speckle activity. Additional GSEA and protein complex analysis of *MALAT1* enriched proteins confirmed significant enrichment of the spliceosomal complex and nuclear speckle factors (**Figure 2I, S5I**). *ITS1* ISH-POCA significantly enriched important nucleolar proteins involved in small subunit pre-rRNA processing (NOL6, NOL11, NOP58, and NOP56) and core nucleolar factors FBL and NCL, consistent with *ITS1*’s nucleolar localization. *ITS1* ISH-POCA also enriched polymerase subunits POLR1A, B, and E which function in nucleolar rRNA transcription^92–94^ and the associated TNF-alpha/NF-κB signaling complex 5 (**Figure 2G,H,I, S5I**). Additionally, the small-subunit processome and nucleolus cellular component were enriched by GSEA for *ITS1* ISH-POCA (**Figure S5I**). Eight members of the RNA-dependent DEAD box ATPase (DDX) family were also enriched in *ITS1* ISH-POCA, some of which, such as DDX21^95^ and DDX18^96^, have known nucleolar function. DDX proteins are RNA helicases with a proposed role in organelle phase separation^97,98^. The potential role of DDX50 and DDX24, which were enriched by *ITS1* ISH-POCA, at the nucleolus remains to be seen. In line with plausible nucleolar function, DDX24 mutation has been shown to alter NPM1 behavior in certain disease contexts^99^, and BioPlex^61,62^ analyses show that both DDX24 and DDX50 interact with known nucleolar proteins such as DDX21. Together, the *MALAT1* and *ITS1* data demonstrate that ISH-POCA quantifies ncRNA-proximal proteomes from distinct nuclear regions and confirms canonical protein networks at both speckles and nucleoli.

### IF-POCA maps protein-targeted proximal proteomes of nuclear substructures

Building on ISH-POCA, the first (to our knowledge) DNA- and RNA-targeted photocatalytic proximity labeling with a proteomics readout, and inspired by antibody-based photocatalytic methods^36,39^, we developed a protein-targeted IF-POCA proteomic method to address three overarching goals: (1) to demonstrate simultaneous high-specificity imaging and proteomics for target proteins at diverse subcellular structures, (2) to rigorously compare the antibody- and ISH-based approaches to benchmark the POCA platform for diverse categories of baits, and (3) to reveal functionally important and spatially constrained interactions by leveraging a comparative proteomic approach for both our ISH and IF methodology. We repurposed our IF-POCA imaging workflow (**Figure 1**) for proteomics and targeted two proteins-of-interest matched with the compartments that we analyzed via ISH-POCA: NPM1 (nucleoli) and SON (nuclear speckles). We added NUP98 (nuclear membrane) to our primary antibody panel to further assess the generalizability of the IF-POCA methodology across diverse cellular structures (**Figure 1C,D, 3A**). As with ISH-POCA, imaging confirmed specific targeting of IF-POCA (**Figure 3C**).

**Figure 3.**
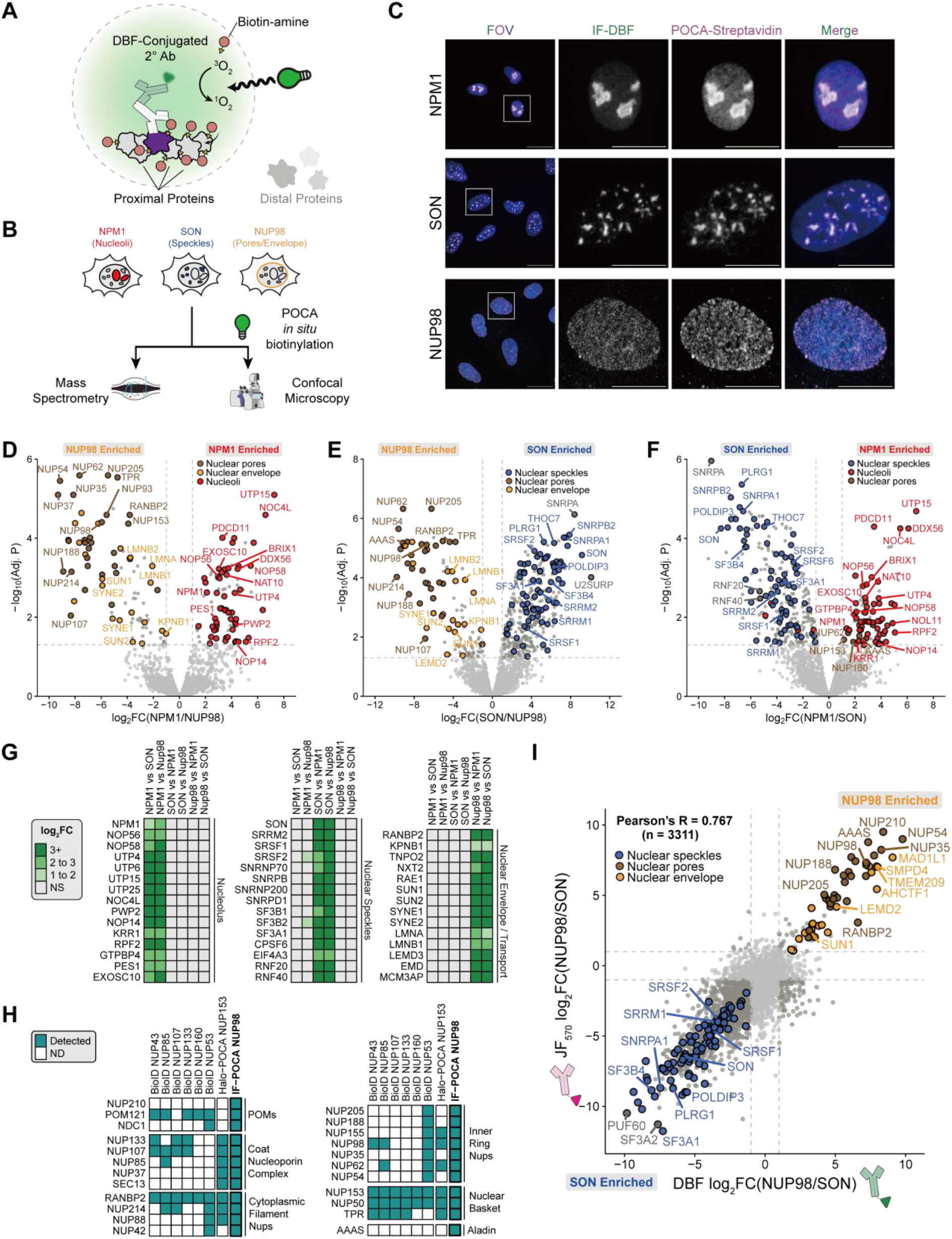
IF-POCA uncovers diverse nuclear substructural proteomes. **A)** Schematic showing the IF-POCA labeling approach. **B)** Schematic showing the experimental workflow for IF-POCA targeting distinct nuclear compartments. **C)** Representative confocal images from each of the three benchmarked IF-POCA targets, depicting IF-DBF (green), POCA-streptavidin (magenta), a merged image of both labeling channels plus DAPI (blue), and a cropped image of a single cell from the merged image. Overlap between DBF and streptavidin signal appears as white. All images are maximum intensity projections. Scale bars are 20 µm in the standard images and 10 µm in the cropped images. **D–F)** Volcano plots of proteins identified in pairwise bait proteins between NPM1, SON, and NUP98. Each dot represents a single protein. Proteins of interest are labeled as follows; red for nucleoli, brown for nuclear pore, orange for nuclear membrane, blue for nuclear speckles. Nucleoli, nuclear speckle, and nuclear envelope/pore annotation lists were generated from the Human Protein Atlas (HPA)^15^, CORUM^80^, and UniProtKB^81^ databases. Hashed lines delineate the thresholds for significance, with greater than 2-fold change in relative enrichment and an FDR-corrected p-value less than 0.05. **G)** Heatmap summarizing log_2_ fold-changes across the three baits versus the no-primary negative control, highlighting enrichment of expected marker proteins for nucleoli, nuclear speckles, and nuclear pore/membrane. **H)** Heatmap of proteins identified by NUP98 IF-POCA and previous studies using BioID^107^ and Halo-POCA^38^. “Detected” indicates at least one spectral count. **I)** Comparison of IF-POCA log_2_ fold-change between SON and NUP98 baits using DBF-versus JF_570_-conjugated secondary antibodies. All raw and processed proteomics data associated with this figure can be found in **Table S6**.

Focusing on the nuclear protein targets, IF-POCA significantly enriched 119 NPM1-proximal, 405 SON-proximal, and 126 NUP98-proximal proteins (**Figure 3D-G, S6A**). Of the significantly enriched proteins, 78 NPM1 hits (65.5%) were nucleolar annotated, 184 SON hits (45.4%) were nuclear speckle annotated, and 66 NUP98 hits (52.4%) were nuclear pore or nuclear envelope annotated in at least one of the Human Protein Atlas (HPA)^15^, CORUM^80^, and UniProtKB^81^. NPM1 IF-POCA enriched factors across successive stages of ribosome biogenesis (**Figure 3D,F-G, S6B**), from early 2’-O-methylation of pre-rRNA by Box C/D snoRNP components (NOP56, NOP58) through 18S rRNA processing mediated by core SSU processome subunits of the UtpA (UTP4, UTP15) and UtpB (PWP2, UTP25) sub-complexes and cleavage factors NOP14 and KRR1 (**Figure S6B,C**). Later-acting factors such as NOC4L, PES1, RPF2, and GTPBP4 further mark pre-40S maturation, 5S RNP integration onto the pre-60S particle, and large subunit export. The nuclear RNA exosome subunit EXOSC10 was recovered, consistent with its role in nucleolar pre-rRNA quality control^102,103^. Beyond the nucleolus, NPM1 IF-POCA enriched nuclear pore complex proteins NUP153, NUP160, NUP62, and AAAS relative to SON IF-POCA, likely reflecting the peripheral positioning of nucleoli against the nuclear envelope and NPM1’s established role as a nucleocytoplasmic shuttle^104–106^.

NUP98 IF-POCA enriched 126 proteins spanning the nuclear pore complex, transport machinery, nuclear envelope, and lamina, including all 33 human nucleoporins—from membrane-embedded POMs to the nuclear basket and cytoplasmic filaments—representing a substantial improvement in coverage over previous nuclear pore proximal datasets (**Figure 3D-E,G-H, S6D**)^38,107108^. Enriched transport factors reflect the bidirectional trafficking through NUP98’s FG repeat-rich channel, capturing major importins (KPNB1, KPNA2), exporters (XPO1, CSE1L), and mRNA transport factors (NXT2, TNPO2). Two functionally interesting hits anchor the transport picture: RAE1, which binds NUP98 to form a conserved mRNA export complex that hands mRNPs to pores, and MCM3AP (GANP), the scaffold of the REX-2 complex that coordinates NXF1-dependent mRNA shuttling to pores^109–111^. The NUP98-proximal proteome then extends outward beyond the pore itself into the nuclear envelope and lamina, enriching the full LINC complex (SUN1, SUN2, SYNE1, and SYNE2), the lamins LMNA and LMNB1, and the LEM-domain proteins EMD and LEMD3.

SON IF-POCA significantly enriched an interconnected set of splicing proteins that canonically define nuclear speckles, including the U snRNP components (U1-U5), the full SF3a and SF3b complexes of U2 snRNP, and Sm ring proteins, as well as SRSF1, SRSF2 (SC35), and SRRM2 (**Figure 3E-G, S6B,C**). SON IF-POCA enriched multiple steps of speckle-associated co-transcriptional processing from splice site recognition through transcript maturation and export. This included enrichment of pre-mRNA 3’-end processing factors (11 enriched), catalytic activation of the spliceosome by the Prp19 complex^112^, EJC components (6 enriched) and the transcriptional handoff machinery of the THO/TREX complex^113–117^. The m6A writer component VIRMA and the m6A demethylases ALKBH5 and FTO were all enriched, consistent with evidence that the m6A methyltransferase complex is recruited to nuclear speckles^118–121^. The U1 snRNP-binding cohesin ring was also enriched, consistent with cohesin directly regulating alternative splicing at speckle-associated chromatin domains^122,123^. Interestingly, the intact MRN complex (MRE11, RAD50, NBN), the primary sensor of DNA double-strand breaks, appeared unexpectedly, raising the possibility that transcription-coupled DNA damage surveillance is spatially coordinated with splicing machinery at the speckle-chromatin interface^124^.

### POCA is modular and generalizable across subcellular compartments

Two key design principles of POCA are the modular, interchangeable photosensitizer, and target flexibility. While we demonstrated that POCA could readily be performed with DBF, we also found that POCA could be readily adapted to use JF_570_, or commercially available dyes ATTO Thio12 and Eosin 5 (**Figure 3I, S1B-D, S7A-C**). Notably, for paired immunofluorescence, Eosin 5 and ATTO Thio12 showed dimmer FISH signals (**Figure S7A,B**). As expected, fluorophores with high fluorescence quantum yield and low singlet oxygen quantum yield (ATTO 488, ATTO 565) did not show any evidence of biotin deposition (**Figure S7A,B**). To demonstrate the modularity of POCA to define proximal proteomes, we repeated the IF-POCA experiments targeting NUP98 and SON using JF_570_ as the photosensitizer instead of DBF. JF_570_ and DBF IF-POCA generated highly consistent IF and fluorescent streptavidin signals as well as proximal proteome measurements (**Figure 3I, S7C**), emphasizing the interchangeability of photosensitizers within POCA workflows. Highlighting POCA’s consistency, recently validated nuclear speckle proteins DHX38, RBM15, SAP18, PQBP1, and PPIL4 were enriched with both JF_570_ and DBF IF-POCA targeting nuclear speckles^82^.

In addition to the subnuclear compartment analysis, we used IF-POCA to target CENP-A (centromere), COXIV (inner mitochondrial membrane), and TOM20 (outer mitochondrial membrane) to further assess the generalizability of IF-POCA across diverse cellular structures (**Figure 1C,D, S3D, S8A-H**). We compared IF-POCA experiments for CENP-A versus NUP98 and COXIV versus TOM20, to investigate POCA’s compatibility with additional nuclear and extranuclear targets (**Figure S8A-H**). Notably, while the inner and outer membranes of mitochondria are generally only 20 nm apart^125^, POCA resolved significant differential proximal proteomes for the inner mitochondrial membrane protein COXIV and outer mitochondrial membrane TOM20 (**Figure S8A-F**). Comparative analysis with confirmed COXIV proximal proteins were enriched for inner mitochondrial proteins, including oxidative phosphorylation complex component proteins, while TOM20 proximal proteins were enriched for mitochondrial outer membrane proteins, including those involved in mitochondrial fission and fusion (**Figure S8A-F**). Together, these data demonstrate that IF-POCA can robustly capture proximal proteomes at distinct subcellular and suborganellar structures.

### Shared ISH-POCA-and IF-POCA-enriched subnuclear proteins define high confidence proximal proteomes for nuclear speckles and nucleoli

The modality-agnostic POCA labeling chemistry affords a unique opportunity to directly compare the proximal proteomes of targeted proteins and nucleic acids. As such, we compared the proteins enriched in our matched ISH-POCA and IF-POCA datasets at nucleoli and nuclear speckles (**Figure 4A**). We reasoned that these analyses would provide an additional layer of stringent benchmarking for the POCA platform, analogous to the added confirmation provided by a reciprocal co-immunoprecipitation when compared to single bait^126^. Though they target inherently different biomolecules, ISH-POCA and IF-POCA proteome enrichment targeting an RNA or protein, respectively, in a shared subcellular space were significantly correlated (**Figure 4B**). The intersection of ISH-POCA and IF-POCA datasets converged on highly specific components of each nuclear body, yielding subnuclear proteomes strongly enriched for core nucleolar and nuclear speckle proteins (**Figure 4C–E, S9A**). Overall, 100% (35 of 35) of the shared proteins between NPM1 IF-POCA and *ITS1* ISH-POCA were annotated to have nucleolar localization (**Figure 4D**)^15,80,81^. Additionally, 80% (69 of 86) of the shared proteins between *MALAT1* ISH-POCA and SON IF-POCA had known nuclear speckle localization (**Figure 4E)**^15,80,81^.

**Figure 4.**
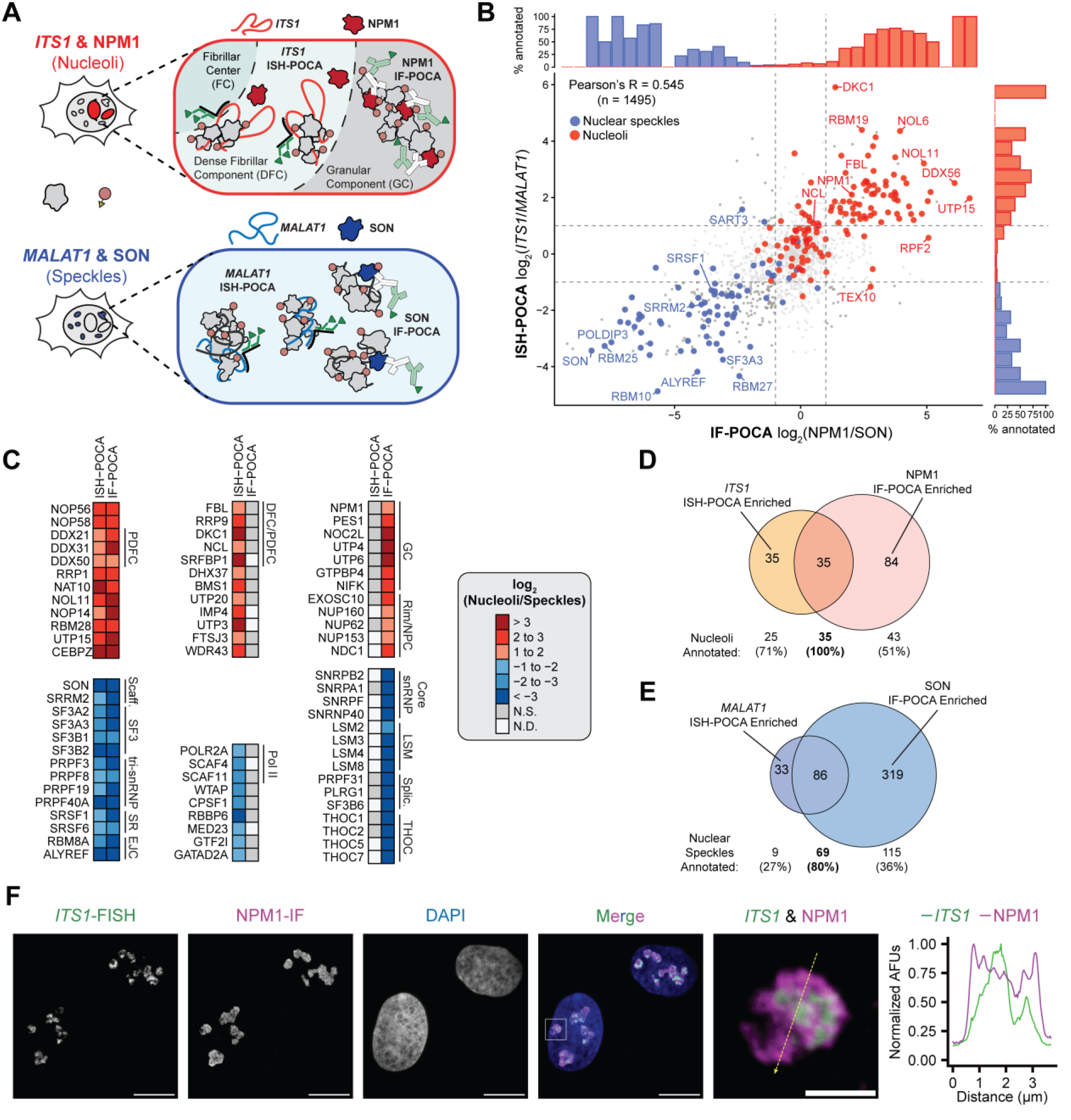
Complementary POCA labeling methods illuminate differential nuclear substructural proteomes through multiple viewpoints. **A)** Schematic depicting IF-POCA and ISH-POCA labeling approaches for SON and *MALAT1* targeting of nuclear speckles, NPM1 targeting of nucleolar GC, and *ITS1* targeting of nucleolar FC. **B)** Enrichment distribution of known nucleolar and nuclear speckle proteins as defined by the HPA^15^. Dashed lines denote the threshold of enrichment, log_2_ fold-change > 2. Histograms depict the fraction of proteins in a particular range of log_2_ fold-change of width 0.5 that have annotations for nuclear speckles (blue) or nucleoli (red) for each method, expressed as a percentage. **C)** Heatmap summarizing log_2_ fold-changes of expected marker proteins for nucleoli and nuclear speckles across IF-POCA and ISH-POCA labeling of unique baits within the same nuclear subcompartment. **D & E)** Venn diagrams of the overlap between *ITS1* ISH-POCA and NPM1 IF-POCA nucleolar proteomes (**D**) and between *MALAT1* ISH-POCA and SON IF-POCA nuclear speckle proteomes (**E**), with nucleolar- and speckle-annotated protein counts indicated below each region, where annotation lists were generated from the Human Protein Atlas (HPA)^15^, CORUM^80^, and UniProtKB^81^ databases. **F)** Representative images of single optical sections depicting *ITS1* FISH (green) and NPM1 IF (magenta) co-FISH imaged with a SoRa CSU-W1 super-resolution spinning disk microscope. A merged image of both labeling channels with DAPI (blue), and a cropped image of a signal pattern from the merged image. Overlap between *ITS1* FISH (green) and NPM1 IF (magenta) appears white. Scale bars are 10 µm in the standard images and 2 µm in the cropped image. The line scan graph shows fluorescence intensity of *ITS1* FISH (green) and NPM1 IF (magenta) along the dotted yellow line in the adjacent representative image.

Proteins in the intersection of reciprocal ISH-POCA and IF-POCA datasets were highly specific to annotated nucleolar and nuclear speckle proximal proteomes (**Figure 4C**). Shared proteins between NPM1 IF-POCA and *ITS1* ISH-POCA included well characterized nucleolar proteins such as nucleolin (NCL)^127–129^, RNA helicases DDX21^95^ and DDX18^96^, and ribosomal RNA maturation factors PES1^130,131^ and BOP1^132,133^ (**Figure 4C**). These shared proteins were enriched for complexes critical to proper rRNA maturation in the nucleolus^134,135^, including small nucleolar RNA (snoRNA) binding and the small nucleolar ribonucleoprotein NOP56p-associated pre-rRNA complex (**Figure S9B**).

The shared nuclear speckle proteins between SON IF-POCA and *MALAT1* ISH-POCA included canonical markers SON, SRRM2, and SC-35 (SRSF2) as well as the SF3, tri-snRNP, and EJC complexes (**Figure 4C**). Additionally, CORUM^80^ protein complexes enriched in the shared *MALAT1* and SON datasets included known nuclear speckle protein complexes such as the spliceosome, the 17S U2 snRNP, and the CDC5L complex (**Figure S9B**). Among the non-speckle-annotated proteins enriched by both IF-POCA and ISH-POCA at nuclear speckles, several were consistent with the splicing-related functional logic of nuclear speckle biology. Enrichment of the 3’ end cleavage and polyadenylation (CPA) factors CSTF2 and CPSF2 was consistent with their prior proteomic detection in nuclear speckle biochemical fractions^136^. Enrichment of RBBP6 confirmed the recent finding that this CPA scaffold protein physically anchors polyadenylation machinery to nuclear speckles to facilitate efficient 3’-end processing of highly expressed genes^114^. Similarly, the shared enrichment of FUBP1—as well as related proteins SF1, U2AF2, and SF3B1—suggests it may localize to the nuclear speckle to aid in 3’-splice site recognition^137^. Though FUBP1 was absent from prior speckle proteomic datasets and is best known as a transcriptional activator of c-Myc, localization at speckles further establishes its function at the transcription-processing interface^137^. Finally, ubiquitin-dependent protein turnover is tightly linked to splicing regulation: ubiquitination facilitates spliceosome assembly and modulates splicing factor activity, while active proteasomes concentrate at nuclear speckles where they degrade splicing components and influence speckle organization^138^. Consistent with this, our nuclear speckle proximal proteomes recovered the proteasome activator PSME3, a known structural organizer of nuclear speckles, and the exclusively nuclear E3 ligase CUL4B, suggesting that localized ubiquitin-mediated protein turnover is an active feature of speckle biology^139,140^. These data suggest a route by which active ubiquitin-mediated protein turnover at nuclear speckles enables rapid cycling of the elongation and splicing complexes required to sustain the elevated transcriptional output of speckle-proximal genes^139,141^.

To extend the cross-validation framework to a third nuclear structure, we compared enriched proteomes at centromeres from Pan Alpha ISH-POCA and CENPA IF-POCA proteomics (**Figure S9F**). As with nucleoli and nuclear speckles, the intersection between the two methods provided enhanced selectivity. All 7 shared significantly enriched proteins represented bona fide centromere resident proteins, spanning CENP family members (CENPB, CENPC, CENPU, CENPL, CENPI), the inner centromere scaffold INCENP, and the pericentromeric heterochromatin regulator SMCHD1. Furthermore, individual CENP family members were broadly enriched across both Pan Alpha ISH-POCA and CENPA IF-POCA relative to their respective spatial controls (Telomere and NUP98) (**Figure S9G,H**). Together, the shared ISH-POCA and IF-POCA datasets demonstrate that targeting a protein and a nucleic acid at the same structure through independent POCA platform approaches provides built-in cross-validation, establishing a framework for high-confidence subcellular proteome definition.

### Defining distinct proximal proteomes of nuclear substructures through paired ISH-POCA and IF-POCA

In addition to shared proteomes, we postulated that our spatially-matched IF-POCA and ISH-POCA analyses could identify sub-compartment differences in protein composition surrounding target RNAs or proteins. These comparisons take advantage of POCA’s enhanced resolution for spatially restricted labeling, e.g., mitochondrial inner versus outer membrane labeling (**Figure S8E,F**). Consistent with this hypothesis, IF-POCA and ISH-POCA identified distinct sets of nuclear speckle and nucleolar proteins.

At nuclear speckles, the 33 proteins uniquely enriched by *MALAT1* ISH-POCA were significantly (FDR-adjusted P < 0.05) overrepresented for GO annotations associated with RNA splicing and nuclear speckle localization (**Figure S9C**). Notably, *MALAT1* ISH-POCA uniquely captured transcriptional machinery, including Pol II subunit POLR2A and Pol II adapters SCAF4 and SCAF11, supporting *MALAT1*’s localization at transcriptionally active chromatin^142,143^ (**Figure 4C**). IF-POCA captured 319 proteins unique to SON (**Figure 4E**). Proximal to SON we observed a broader range of proteins involved in RNA-mediated processes, including spliceosomal components, snRNP and LSm proteins, the hnRNP family of splicing regulators, polyadenylation and 3’ end processing proteins, and the THOC complex (**Figure 4C**). These proximal proteins are highly consistent with SON’s established scaffolding function throughout the interior of nuclear speckles^84,85,144,145^.

Mammalian interphase nucleoli are organized into phase-separated subcompartments with distinct proteomic compositions. To test whether POCA could resolve these sub-nucleolar features, we stratified the protein subsets unique to the *ITS1* ISH-POCA and NPM1 IF-POCA nucleolar datasets (**Figure 4A**). Nucleoli are multilayered organelles made up of a fibrillar center within a dense fibrillar component which is further surrounded by a granular component^146,147^. The dense fibrillar component is a hub for rRNA processing which contains a fibrillar center for rRNA transcription^146,147^. The granular component surrounds the dense fibrillar component and mediates genome-nucleolus interactions and the final stages of nucleolar ribosome processing^146,147^. By smFISH analyses, *ITS1* localizes predominantly to the dense fibrillar component, while NPM1 localizes predominantly in the granular component, where it helps define the nucleolar border and pre-ribosome assembly occurs^148,1494^. High-resolution co-FISH imaging confirmed overlapping localization of *ITS1* and NPM1 in hTERT-RPE1 cells (**Figure 4F**). NPM1 staining partially overlapped with *ITS1* FISH signal but extended beyond it, consistent with NPM1 localizing to both the dense fibrillar component and the surrounding granular component (**Figure 4F**)^150^. Accordingly, *ITS1* ISH-POCA and NPM1 IF-POCA enriched distinct but overlapping sets of canonical nucleolar factors (**Figure 4C**).

Beyond the overlapping region, we observed that *ITS1* ISH-POCA alone significantly enriched the dense fibrillar component markers nucleolin (NCL) and dyskerin (DKC1) (**Figure 4C, S9D**)^151^. This enrichment pattern aligns with the expected *ITS1* sub-nucleolar localization and is consistent with POCA labeling occurring with subnucleolar spatial precision. Similarly, NPM1 IF-POCA uniquely captured 84 proteins that were not significantly enriched by *ITS1* ISH-POCA (**Figure 4C,D**). We confirmed that 11 of these proteins had imaging-validated granular component localization, which corroborated that POCA could achieve specific proximity labeling with sub-nucleolar resolution (**Figure S9D**)^151^. Among the remaining proteins uniquely enriched with NPM1 IF-POCA we observed several membrane-embedded and nucleoplasmic-facing nucleoporins (NUP153, NUP160, NUP62, NDC1, **Figure 4C**). NUP153 in particular is critical to facilitating pre-ribosomal subunit nuclear export, but colocalization between NPM1 at the nucleolar rim and the nuclear basket has not previously been shown^152,153^. The selective recovery of these nucleoporins by NPM1 IF-POCA and not *ITS1* ISH-POCA likely reflects the spatial proximity of nucleoli with nuclear pores during pre-ribosome export and captures the precision of POCA to resolve subnuclear structure contact sites.

## Discussion

We demonstrate that the POCA platform is a modular photocatalytic proximity labeling platform using a single photosensitizer that can covalently label DNA-, RNA-, and protein-proximal biomolecules. POCA can leverage established (DBF), new (JF_570_), and commercially available photosensitizers (ATTO Thio12, Eosin 5). These can be coupled with alternative proximity labeling chemistry for both direct and click chemistry-based covalent labeling. Importantly, POCA’s recovery efficiency of proximal proteomes enables DNA, RNA, or protein targeting with as few as 2 million cells per sample replicate.

POCA achieves more precise, spatially restricted, and sensitive covalent labeling compared to peroxidase-mediated proximity labeling by limiting diffusion of reactive oxygen species, consistent with previous photoactivated proximity labeling approaches^36,39,154^. By balancing fluorescence for targeting validation and singlet oxygen generation for biotin deposition, ISH-POCA obtained more specific protein enrichment at telomeres and pericentromeres compared to our recent peroxidase-based datasets^32^. Based on our work and others, ISH-directed proximity labeling can target unique RNAs, repetitive genomic loci, single copy loci, and even haplotype resolved loci^30–32^. We believe the improved proximity labeling with ISH-POCA should also enable targeting of small genomic regions, allowing future ISH proximity labeling work to push to smaller targets than our recent HRP-based targeting of 80-kb single-copy *HOX* gene clusters^32^.

Our ability to apply POCA to target different biomolecules within the same nuclear compartments enabled us to investigate nucleolar (NPM1 and *ITS1*), nuclear speckle (SON and *MALAT1*), and nuclear membrane (NUP98) proximal proteomes. These experiments complement recent efforts to measure proximal proteomes of discrete nuclear bodies using TurboID cell lines^82,155^, although SON and NUP98 were not targeted in these studies. Notably, both TurboID and POCA identified regulatory proteins such as the E3 ubiquitin ligase RNF20 as enriched at nuclear speckles. RNF20 was enriched at speckles through RBM15B targeted TurboID and SON/*MALAT1* targeted POCA. Given these results, future lines of inquiry could target other constitutive proteins to map membranous and phase-separated subcellular compartments (e.g., synapses, Cajal bodies, PML bodies, paraspeckles). Because POCA targeting does not require genetic manipulation of cell lines, extensions of these nuclear structure proteome studies using POCA would simplify analyses of proximal proteomes in different mutational contexts, such as leukemia-associated NUP98 rearrangements and onco-fusions which could help gain key insights into proteome-level misregulation in cancer^156,157^.

Previous studies demonstrated that enzyme-based proximity labeling methods such as TurboID and APEX can be deployed at multiple subcellular structures to measure protein shuttling within the cell^158^. By combining IF-POCA and ISH-POCA, we extend this principle to distinct biomolecular classes. Protein and nucleic acid targets can be used as independent anchors for complementary proximity profiling within a unified experimental framework. Importantly, labeling remains compartment-specific, *MALAT1* ISH-POCA at nuclear speckles did not recover proteins enriched by NPM1 IF-POCA at nucleoli, and vice versa. These results confirm that each targeting modality interrogates a spatially discrete microenvironment. Furthermore, the resolving power of this combined approach is well illustrated within the nucleolus itself. *ITS1* ISH-POCA preferentially enriched factors associated with the fibrillar center and dense fibrillar component, whereas NPM1 IF-POCA captured proteins of the granular component and nucleolar rim, consistent with the established subcompartmental localization of these two molecules^148,149^. These results demonstrate that POCA can distinguish adjacent subnuclear domains that are organized by both RNA and protein scaffolds, which has broad implications for studying compartments whose architecture depends on the interplay between these two biomolecular classes. More broadly, the ability to anchor proximity labeling to either a nucleic acid or protein target within the same platform creates new opportunities to investigate nucleocytoplasmic transport, stress-dependent protein relocalization, and disease-associated changes in compartment composition^159–161^. For example, combined ISH-and IF-POCA could be used to compare the proximal proteome of nucleolar *ITS1* with that of NPM1 mutants that mislocalize to the cytoplasm in acute myeloid leukemia^160^, disentangling RNA-anchored from protein-anchored changes in nucleolar organization in disease.

While the current POCA platform was successfully implemented in a variety of cell lines and targeting diverse molecular environments, there are important limitations to consider during experimental design. Formaldehyde fixation can induce structural changes to compartment architecture^162^ and artifactually modify proteins that will later be labeled and purified. Thus, complementary studies using alternative fixation strategies may prove valuable in certain contexts. Low abundance proteins or those with diverse post-translational modifications may also pose a challenge for detection of proximal proteins by mass spectrometry-based proteomics. Fixation also prevents the current implementation of POCA from being applied to live cells. ISH probes can feasibly cover almost the entire human genome^163,164^, though certain RNAs and genomic regions, such as low abundance RNAs, may be hard to target with ISH-POCA without higher cell input. Similar to canonical IF, the spatial precision of IF-POCA is limited to the specificity of the chosen primary antibody. POCA largely labels histidine residues^38^, therefore it stands to reason that our labeling approach may be biased for proteins with surface exposed histidine residues, as has been characterized for other proximity labeling approaches^165^. Therefore, future iterations of POCA proximity labeling may benefit from alternative labeling chemistry that enables for more promiscuous residue labeling with biotin^44^.

In conclusion, we present a modular photocatalytic proximity labeling platform capable of mapping the local proteomes of DNA loci, RNAs, and proteins. POCA labeling does not require genetic engineering and can use oligos and antibodies conjugated with commercially available photosensitizers. Using relatively small numbers of cells, we demonstrate clear spatial precision and adaptability to varied targets in multiple cell lines using standard ISH and IF workflows. Given the targeting flexibility of the POCA platform, we envision its widespread adoption to produce high-confidence interactome maps of nucleic acid and protein proximal protein networks across diverse cellular contexts.

## Methods

### Cell culture

hTERT-RPE1 human epithelial cells (gift from Joshua Vaughan), U2-OS human epithelial cells, and EY.T4 mouse fibroblasts were cultured in ATCC-formulated DMEM (Gibco, 11965-092) supplemented with 10% fetal bovine serum (FBS) and 100 U/mL penicillin–streptomycin. K562 human myelogenous leukemia cells were grown in ATCC-formulated RPMI (Gibco 11875-093) supplemented with 1x non-essential amino acids (Fisher Scientific, 11-140-050) and 1 mM sodium pyruvate (Fisher Scientific, 11-360-070). All media was filtered (0.22 µm) before use. All cells were grown at 37 °C in a humidified atmosphere containing 5% CO_2_. Cell lines were tested monthly for mycoplasma using the mycoplasma detection kit (InvivoGen).

### Primary Oligo Probes

Primary oligos targeting the human alpha satellite repeats and telomeres, and ITS RNA were purchased as individually column-synthesized DNA oligos from Integrated DNA Technologies. Primary oligos targeting human MALAT1 RNA were designed using PaintSHOP and ordered in oPool format from Integrated DNA Technologies.

### Primer Exchange Reaction (PER)

To extend primary oligos into PER concatemers, 100-µL reactions were prepared containing 1× PBS, 10 mM MgSO_4_, 300 µM dNTP mix (dA, dC and dT only), 100 nM Clean.G hairpin, 50 U/mL Bst LF polymerase (New England Biolabs), 500 nM hairpin and water to 90 µL. Reactions were incubated for 15 min at 37°C, and then 10 µL of 10 µM primary oligo probe was added to each reaction, which were incubated at 37 °C for 2 h, followed by 20 min at 80°C to heat inactivate the polymerase. To assess extension efficiency and concatemer lengths, samples (diluted in 2× Novex TBE-Urea Sample Buffer, Thermo Fisher Scientific LC6876) and DNA ladders (Low Range and 100 bp or 1 kb Plus, Thermo Fisher Scientific) were boiled at 95°C for 5 min and then were run on pre-heated 10% TBE-urea gels (Thermo Fisher Scientific EC68755BOX) for 45 min at 70°C using a circulating water bath connected to the gel cassette. Reactions were dehydrated via vacuum evaporation at 60°C then stored at -20°C until hybridization.

### Amine-Modified Oligonucleotide-Photosensitizer Bioconjugation

Amine-modified oligonucleotides were purchased from Integrated DNA Technologies (Newark, NJ) and reconstituted in 100 mM sodium tetraborate decahydrate pH 8.5 to a concentration of 1 mM. The amine-modified p27*p27* oligonucleotide was ordered as follows: /5AmMC6/TTTATGATGATGTATGATGATGT. To a 1.5-mL microcentrifuge tube was added 20 µL (20 nmol) of amine-modified oligonucleotide, 16 µL (400 nmol, 20 eq.) of freshly prepared 25 mM DBF-NHS or JF_570_-NHS in DMSO, and 64 µL of 100 mM sodium tetraborate decahydrate pH 8.5, and the reaction was incubated at room temperature for 18 h, protected from light. Then, to the reaction mixture was added 200 µL of UltraPure water, 20 µL (1/10 volume) of 3 M NaCl (aq.), and 500 µL (2.5 volumes) of ice-cold ethanol, immediately vortexing at maximum speed for 30 s to initiate oligonucleotide precipitation. The reaction mixture was then frozen at -80 °C for 2 h. Oligonucleotide conjugates were pelleted by centrifugation at 20,000 g for 30 min at 4 °C, rinsed with 500 µL of ice-cold 70% ethanol, taking care not to disturb the pellet, and centrifuged at 20,000 g for 15 min at 4 °C. After removing the supernatants, the oligonucleotide conjugate pellets were dried at room temperature for 15 min. Each pellet was then resuspended in 50 µL of UltraPure water and purified by reverse-phase semi-preparative HPLC (Agilent 1260 Infinity equipped with an Agilent InfinityLab Poroshell 120 EC-C18, 4.6 x 150 mm, 4 μm column) using a 30-minute linear gradient from 0–25% acetonitrile in 100 mM triethylammonium acetate pH 8. Conjugate-containing fractions were analyzed by native PAGE gel (Invitrogen EC63155BOX) to confirm DBF conjugation and purity, then subsequently lyophilized and stored at -20 °C until further use.

### Fixation for ISH-POCA

For each ISH-POCA experiment, 300,000 cells were seeded into each well of a glass-bottom 6-well plate (Cellvis P06-1.5H-N) to culture for 24 hours to reach 750,000 - 1 million cells per well. Cells were briefly rinsed once with DPBS then fixed in 4% paraformaldehyde (*w/v*) (Electron Microscopy Sciences 15710) in PBS at room temperature for 10 minutes. Fixed cells were then washed with DPBS three times for 5 minutes each. Fixed cells were stored in fresh DPBS at 4 °C until solid-phase hybridization. Fixed cells were used within 5 days. For K562 cells, each well of the 6-well plate received 0.1 mg/mL poly-D-lysine (Fisher Scientific A3890401) and incubated on a shaker for 1 hour followed by three rinses with 1x PBS. 2 million cells were then plated per well in growth media and placed in a 37 °C CO_2_ incubator for 30 minutes. The media was carefully removed with a pipette and the cells were carefully fixed in 4% PFA for 10 minutes followed by three washes with 1x PBS.

### Oligo hybridization in fixed cells

Oligo DNA hybridizations were performed on cells fixed to glass, with all washes being performed in 1 mL. After fixing, cells were rinsed once with fresh phosphate buffered saline (PBS), and permeabilized in PBS-0.5% TritonX-100 (Sigma T8787) for 10 minutes. After a PBS-0.1% Tween 20 (PBS-T) (Sigma T2287) wash, permeabilized cells were incubated in 0.1 N hydrochloric acid (HCl) for 5 minutes. After two washes for 2 minutes each in 2x saline sodium citrate-0.1% Tween20 (2X SSC-T), cells were incubated in 2X SSC-T-50% formamide for 20 minutes on a heat block in a 60 °C water bath. Cells were exchanged into 500 µL per well of primary hybridization buffer comprised of 2X SSC-T, 50% (vol/vol) formamide, 10% (*w/v*) dextran sulfate, 0.4 μg/ul RNAse A, ∼500 nM extended primary oligos, 0.1% Tween 20, and UltraPure water. The cells were denatured on a heat block in a water bath at 78 °C for 3 minutes. The cells were then hybridized with primary oligos at 37 °C overnight with nutation for approximately 18 hours. The next day, cells were washed with pre-heated 60°C 2X SSC-T four times for 5 minutes each followed by two 2X SSC-T washes for 2 minutes each at room temperature. Fully washed cells were then incubated in 100 nM secondary DBF oligo that hybridizes to the concatemer sequence on the primary oligo diluted in PBS for 1 hour at 37 °C. Cells then underwent three 5-minute washes in PBS-T at 37 °C and one PBS wash at room temperature.

Oligo RNA hybridizations were performed in the same manner as DNA hybridization with a few changes. While preparing the cells for primary hybridization the cells were <u>not</u> incubated in 0.1 N HCl. Hyb1 buffer for RNA targets was formulated <u>without</u> RNAse A, this volume was replaced with UltraPure water. Cells were incubated at 60 °C for 3 minutes instead of 78 °C, prior to overnight incubation at 37 °C.

### ISH-POCA Biotin Labeling

After the post-secondary hybridization washes, the cells were incubated for 10 minutes in 1 mL of 2 mM Biotin-PEG3-Amine (Lumiprobe Corporation 2623) diluted in PBS. The samples were then exposed to 508 nm light from an LED (Thorlabs M505L5) with a condenser lens and diffuser (Thorlabs ACL2520U-DG15-A) supplied with 0.35 Å of current while about 10 cm from the light to achieve a power density of approximately 0.73 mW/cm^2^ at the cells in a 6-well plate. Exposure was carried out for 5 minutes to induce biotin labeling through DBF reactivity. The cells were then washed 3x in PBS-T at 37 °C for 5 minutes each.

### Microscopy-based quality control for hybridization and biotinylation of ISH-POCA samples

We sample cells prior to preparing loci specific AP-MS samples to monitor the locus specificity of primary-secondary oligo hybridization and biotin labeling. To assess the hybridization and labeling quality, we included one well of a 6-well plate for each target as well as a non-targeted control. After the post-biotin labeling wash steps, the quality control wells were incubated with 0.5 μg/mL Alexa Fluor 647-streptavidin (Thermo Fisher S32357) in PBS-T, 1% bovine serum albumin at 37 °C for 30 minutes. Stained cells underwent three washes for 5 minutes each in 1 mL PBS-T at 37 °C to remove unbound Alexa Fluor 647-streptavidin conjugate. Washed cells were then incubated in 5 μg/mL DAPI for 15 minutes in the dark. Cells were then washed once in PBS-T and then placed into Slowfade Gold Antifade Mountant for confocal imaging of Alexa-Fluor 647-streptavidin and DBF signals.

### Confocal microscopy for ISH-POCA samples

Confocal imaging was performed using a Yokogawa CSU-W1 SoRa spinning disc confocal device attached to a Nikon ECLIPSE Ti2 microscope. Excitation light was emitted at 30% of maximal intensity from 405 nm, 488 nm, 561 nm, or 640 nm lasers housed inside of a Nikon LU-NF laser unit. Laser excitation was delivered via a single-mode optical fiber into the CSU-W1 SoRa unit. Excitation light was directed through a microlens array disk and a SoRa spinning disk containing 50 μm pinholes to the rear aperture of a 100x N.A. 1.49 Apo TIRF oil immersion objective lens by a prism in the base of Ti2. Emission light was collected by the same objective and directed by a prism in the base of Ti2 back into the SoRA unit, where it was relayed by a 1x lens (conventional imaging) or 2.8x lens (super-resolution imaging) through the pinhole disk and then directed to the emission path by a quad-band dichroic mirror (Semrock Di01-T405/488/568/647-13X15X0.5). Emission light was then spectrally filtered by one of four single-bandpass filters (DAPI:Chroma ET455/50M; ATTO488: Chroma ET525/36M; ATTO565:Chroma ET605/50M; Alexa Fluor 647: Chroma ET705/72M) and focused by a 1x relay lens onto an Andor Sona 4.2B-11 camera with a physical pixel size of 11 um, resulting in an effective resolution of 110 nm (conventional), or 39.3 nm (super-resolution). The Sona was operated in 16-bit mode with rolling shutter readout and exposure times of 70-300 ms.

### Affinity Purification and sample preparation for ISH-POCA proteomics

Biotinylated cells were removed from 4 °C storage to equilibrate to room temperature. 0.4 mL of lysis buffer consisting of 1% SDS and 200 mM EPPS with protease inhibitors (Roche 11836170001) was then added to each well. Lysed cells were then scraped off of the glass and collected in 1.5 mL tubes. The cell mixture was boiled at 95 °C for 30 minutes. The boiled cell mixture was sonicated at 4 °C using a Diagenode Bioruptor Plus at high power for 10 cycles of 30 seconds on and 30 seconds off. The sonicated cell mixture was boiled for a second time at 95 °C for 30 minutes. The boiled lysates were cleared by centrifuging at 21130 G for 30 minutes in an Eppendorf 5424 Microcentrifuge at room temperature. The supernatants were transferred to a fresh 1.5-mL tube. To prevent any remnants of cell debris, the supernatants were cleared for a second time by centrifuging at 21130 G for 30 minutes and the supernatants were transferred to a fresh 1.5-mL tube.

The cleared cell lysates were quantified using the Pierce BCA Protein Assay Kit (Thermo Fisher 23225). Pierce Streptavidin Magnetic Beads (Thermo Fisher 88817) were washed using 1% SDS, 200 mM EPPS lysis buffer three times before use. An equivalent mass of protein from each labeled cell pellet was then used to couple with streptavidin beads in a Protein Lo-Bind tube (Eppendorf EP022431081). The lysates were incubated with the bead slurry for one hour at room temperature with nutation allowing biotinylated proteins to bind. The coupled beads were collected and separated from the flow-through using a magnetic rack (Sergi Lab Supplies 1005a). After the flow-through was removed, the beads underwent the following washes: 1% SDS with 20 mM EPPS twice, 0.1 M Na_2_CO_3_, 2 M urea, and 1 M KCl with 20 mM EPPS twice. All washes were performed as follows: after immobilizing the beads on a magnetic rack for 5 minutes, the supernatant was removed, and the beads were resuspended in the new wash buffer and incubated for 5 minutes with nutation. Finally, the beads were rinsed once with 20 mM EPPS to remove the excess salt.

The washed streptavidin beads were resuspended in 50 μL of 5 mM TCEP, 200 mM EPPS, pH 8.5 for a 20-minute on-bead protein reduction. The proteins were alkylated on-bead using 10 mM iodoacetamide for one hour in the dark. Then DTT was added to the final concentration of 5 mM to quench the alkylation for 15 minutes. The beads were rinsed twice with 200 mM EPPS for on-bead digest. Assuming about 2 μg of eluate protein, 20 ng LysC (Wako) was added to the beads in a 50-μL volume and incubated for 16 hours with vortexing. The next day, 20 ng of trypsin (Promega V5113) was added to the beads and incubated for six hours at 37 °C at 200 rpm. After digestion, the peptide-containing supernatant was collected in a fresh 0.5-mL Protein Lo-Bind tube. Peptides were desalted via the stop and go extraction (StageTip) method and dried in a vacuum concentrator.

### ISH-POCA Sample Mass Spectrometry Data Acquisition

Samples were resuspended in 5% acetonitrile plus 2% formic acid in 200 mM EPPS pH 8.5 (Thermo Fisher, J61476.AK) prior to being loaded onto a Aurora Gen 3 C18 column (IonOpticks, 15 cm, 75 μm I.D.). Peptides were eluted over 90 min gradients running from 98% buffer A (5% acetonitrile, 0.125% formic acid) and 2% buffer B (95% acetonitrile, 0.125% formic acid) to 75% buffer A and 25% buffer B. Sample eluate was electrosprayed (2700 V) into a Thermo Scientific Orbitrap Eclipse mass spectrometer for analysis. High field asymmetric waveform ion mobility spectrometry (FAIMS) was set at “standard” resolution, 4.6 L/min gas flow, and 3 CVs: −40/–60/–80 were used. MS1 scans were conducted at 120,000 resolving power with a 50 ms max injection time, and the AGC target set to 200%. Peaks from the MS1 scans were filtered by intensity (minimum intensity >5 × 10^3^), charge state (2 ≤ z ≤ 6), and detection of a monoisotopic mass (monoisotopic precursor selection, MIPS). Dynamic exclusion was used, with a duration of 90 seconds, repeat count of 1, mass tolerance of 10 ppm, and the “exclude isotopes” option checked. Data-dependent MS/MS scans were performed based on a 1 second cycle time. MS/MS scans were conducted in the linear ion trap with the “rapid” scan rate, 35 ms max injection time, AGC target set to 250%, HCD collision energy of 30%, and 0.7 m/z isolation window.

### Antibody-Photosensitizer Bioconjugation

Goat anti-rabbit IgG (500 µg, Invitrogen A16098) or goat anti-mouse IgG (500 µg, Abcam ab6708) was buffer exchanged into 100 mM sodium bicarbonate pH 8.5 using 7 kD MWCO Zeba spin desalting columns (Thermo Scientific 89882), resulting in a final volume of 200 µL. Then, 10 molar equivalents of DBF-NHS or JF_570_-NHS (2 µL of 10 mg/mL in DMSO) were added and the reaction was incubated at room temperature for 2 h, protected from light. The resulting conjugate was purified by 7 kD MWCO Zeba dye and biotin removal spin column (Thermo Scientific, A44296S), and successful antibody-DBF or antibody-JF_570_ conjugation was verified via SDS-PAGE and UV-Vis analysis. The DBF-conjugated or JF_570_-conjugated anti-rabbit or anti-mouse IgG was stored at 4 °C and used within 1 month.

### IF-POCA Immunostaining and Photocatalytic Labeling

For each IF-POCA experiment, 1.5 million cells were seeded into each 60-mm cell culture dish (Bioland Scientific 705001) and cultured for 20 h to reach 90–100% confluency. Cells were briefly rinsed once with DPBS then fixed in 4% paraformaldehyde (*w/v*) in PBS at room temperature for 10 min. Fixed cells were then washed twice with DPBS and quenched with 125 mM glycine in PBS at room temperature for 5 min. Quenched cells were then washed twice with DPBS and permeabilized with 0.1% Triton X-100 in PBS at room temperature for 10 min. Permeabilized cells were then washed twice with DPBS and blocked with 3% BlockAid (Invitrogen B10710) in PBS at room temperature for 1 h. Blocking solution was aspirated and discarded, and cells were incubated in primary antibodies (**Table S3**) in 3% BlockAid in 0.1% Tween-20 in PBS (PBST) overnight at 4 °C. Primary antibody solution was aspirated and discarded, cells were washed with PBST (3 x 5 min), then incubated in DBF- or JF_570_-conjugated anti-rabbit IgG (1.5 µg/mL in 3% BlockAid in PBST) or DBF- or JF_570_-conjugated anti-mouse IgG (3.0 µg/mL in 3% BlockAid in PBST) at 37 °C for 1 h in a humidifying chamber, protected from light. Secondary antibody solution was aspirated and discarded and cells were washed with PBST (3 x 5 min).

To initiate IF-POCA proximity labeling, immunostained cells were incubated in 5 mM biotin-PEG3-amine (Lumiprobe Corporation 2623) in PBS at room temperature for 5 min and illuminated with a 455-nm LED when using DBF or a 585-nm LED when using JF_570_ (SunLite PAR30, 15 W) at an illuminance of ∼170,000 lx at room temperature for 5 min. Cells were washed with pre-warmed (37 °C) PBST (3 x 5 min), 500 µL of 1% SDS in PBS was added to each dish, and cells were harvested by scraping into 1.5-mL microcentrifuge tubes. Biotinylated lysates were flash frozen and stored at -80 °C until further sample processing.

### IF-POCA Mass Spectrometry Sample Processing

Lysates were thawed, and crosslinks were reversed by incubating at 95 °C for 30 min with gentle shaking (300 RPM), probe sonicating (Misonix Sonicator S-4000, amplitude 50%, 10 cycles, 1 sec on, 10 sec off), and incubating a second time at 95 °C for 30 min with gentle shaking (300 RPM). Lysates were then clarified by centrifugation at 20,000 g for 5 min, and each 500 µL supernatant was transferred to a 1.5-mL low-bind microcentrifuge tube (Fisher Scientific 02-681-320).

For each sample, 20 μL Sera-Mag SpeedBeads Carboxyl Magnetic Beads, hydrophobic (50 μg/μL, total 1 mg) and 20 μL Sera-Mag SpeedBeads Carboxyl Magnetic Beads, hydrophilic (50 μg/μL, total 1 mg) were mixed and washed with water three times. 40 µL of 1:1 SP3 bead slurry was then transferred to each lysate and incubated at RT for 5 min with shaking (1000 RPM). Protein-bead binding was initiated by adding 800 µL of ethanol and incubating at RT for 10 min with shaking (1000 RPM), then the beads were separated from the supernatant on a magnetic rack (Sergi Lab Supplies), the supernatant discarded, and the beads washed 3x with 500 µL of 80% ethanol in UP water. Proteins were eluted from the SP3 beads with 100 μL of 0.2% SDS in PBS at 37 °C for 30 min with shaking (1000 RPM). The elution was repeated once and combined with the initial eluent. Then, total protein concentration was determined by DC assay (Bio-Rad 5000112), and samples were normalized to 200 µg of protein input by diluting with PBS.

Pierce Streptavidin Agarose beads (Thermo Scientific 20353, 50 µL of suspension per sample) were washed 3x with PBS, resuspended in 500 µL of PBS per sample, 500 µL of the resulting streptavidin-agarose slurry was added to each sample, and the lysates were rotated at room temperature for 2 h with nutation, allowing biotinylated proteins to bind. The beads were then pelleted by centrifugation at 1000 g for 1 min and washed once with 6 M urea and 0.2% SDS in PBS, once with 0.2% SDS in PBS, twice with PBS, and twice with UP water. The beads were then resuspended in 200 µL of 6 M urea in PBS, and cysteines were reduced with DTT (10 µL of a 200 mM solution, final concentration 10 mM) at 65 °C for 15 min with shaking (1000 RPM) and alkylated with iodoacetamide (10 µL of a 400 mM solution, final concentration 20 mM) at 37 °C for 30 min. The samples were then diluted with 400 µL of PBS (to achieve a concentration of approximately 2 M urea), the beads were pelleted by centrifugation at 1000 g for 1 min, and the supernatants removed. The beads were resuspended in a digestion buffer consisting of 2 M urea, 4 mM calcium chloride, and 500 ng of trypsin (Worthington Biochemical LS003740) in PBS, and on-bead digest was performed at 37 °C with shaking (250 RPM) overnight. After digestion, the peptide-containing supernatants were transferred to fresh 1.5-mL low-bind microcentrifuge tubes. Peptides were desalted via Pierce C18 pipette tips (Thermo Scientific 87784), dried in a centrifugal vacuum concentrator, then reconstituted with 5% acetonitrile and 1% formic acid in UltraPure water prior to analysis by LC-MS/MS.

### IF-POCA Sample Mass Spectrometry (LC-MS/MS) Analysis

Samples were loaded onto an 18 cm long, 100 μM inner diameter fused silica capillary packed in-house with bulk C18 reversed phase resin (particle size, 1.9 μm; pore size, 100 Å; Dr. Maisch GmbH). Peptides were eluted over a 70 min gradient running from 99% buffer A (water with 0.1% formic acid) and 1% buffer B (80% acetonitrile in water, 0.1% formic acid) to 5% buffer A and 95% buffer B. Detailed gradient parameters: 3 min at 300 nL/min from 1–8% B, 55 min at 300 nL/min from 8–40% B, 6 min at 300 nL/min from 40–55% B, 2 min at 300 nL/min from 55– 95% B, 4 min hold at 95% B. Sample eluate was electrosprayed (2200 V) into a Thermo Scientific Orbitrap Eclipse mass spectrometer for analysis. Data were acquired using a Data-Dependent Acquisition (DDA) method as follows. MS1 scans were conducted in the Orbitrap at 120,000 resolving power with AGC target 250%. Peaks from the MS1 scans were filtered by intensity (minimum intensity 5.0e4), charge state (2–7), and detection of a monoisotopic mass (monoisotopic precursor selection, MIPS). Dynamic exclusion was used, with a duration of 30 seconds, repeat count of 1, mass tolerance of 10 ppm, and the “exclude isotopes” option checked. Subsequent MS2 scans were conducted in the Orbitrap at 15,000 resolving power, HCD collision energy of 30%, and a 1.6 m/z isolation window.

### Microscopy-Based Quality Control for IF-POCA Immunostaining and Proximity Labeling

During each IF-POCA experiment, an imaging quality control sample was prepared for each bait protein in 24-well plate wells containing 12-mm #1.5 glass coverslips (Thomas Scientific 1217N79). After photocatalytic labeling and washing, quality control wells were incubated with 2 μg/mL streptavidin Alexa Fluor 647 conjugate (Thermo Fisher S32357) in PBS at room temperature for 1 h, protected from light. Stained cells were washed with pre-warmed (37 °C) PBST (5 x 5 min), then incubated in 1 µg/mL DAPI in PBS at room temperature for 15 min, protected from light. Cells were then washed once with PBS and mounted onto glass slides using ProLong Diamond Antifade Mountant (Thermo Scientific P36961) for confocal imaging of streptavidin Alexa Fluor 647, DBF, and DAPI signals.

### Confocal Microscopy for IF-POCA Samples

Confocal imaging was performed using a Zeiss LSM880 confocal microscope. Excitation light was emitted at 2–20% of maximal intensity from 405 nm, 514 nm, or 633 nm lasers and focused using an Zeiss alpha Plan-Apochromat 100x/1.46 Oil DIC M27 Elyra objective. The microscope was operated in 16-bit mode with 4X averaging.

### IF- and ISH-POCA Proteomics Protein Identification and Quantification

Raw data collected by LC–MS/MS were searched with MSFragger^166^ (v4.1) and FragPipe (v22.0) with the appropriate UniProtKB reviewed database (downloaded January 5, 2023). Precursor and fragment mass tolerance was set as 0.3 Da (ISH-POCA) or 20 ppm (IF-POCA). A fully tryptic (KR) search space with up to two missed cleavages was used. Peptide length was set to 7–50 and peptide mass range was set to 500–5,000. The following modifications per peptide were set: cysteine carbamidomethylation (fixed, mass shift 57.02146, maximum occurrences 3), N-terminal acetylation (variable, mass shift 42.0106, maximum occurrences 1), methionine oxidation (variable, mass shift 15.9949, maximum occurrences 3). MSBooster^167^ was enabled. Peptide spectrum match validation was performed using Percolator^168^ to a minimum probability of 50% and protein validation was performed using ProteinProphet to a false discovery rate (FDR) of 1%. The MS1 intensity ratio of protein abundances using label-free quantification (LFQ) was determined using IonQuant^169^ (v1.10.38) with normalization enabled and match between runs (MBR) enabled for ISH-POCA and not IF-POCA and then further processed using FragPipe Analyst^170^ (LFQ, raw intensities, Perseus-type imputation, Benjamini-Hochberg adjusted P-values). The resulting data matrix was exported as a CSV file and used for further analyses.

### Fixed-Cell POCA Modified Peptide Analysis Sample Preparation

Consistent with IF-POCA preparation, 1.5 million cells were seeded into a 60-mm cell culture dish (Bioland Scientific 705001) and cultured for 20 h to reach 90–100% confluency. Cells were briefly rinsed once with DPBS then fixed in 4% paraformaldehyde (*w/v*) in PBS at room temperature for 10 min. Fixed cells were then washed twice with DPBS and quenched with 125 mM glycine in PBS at room temperature for 5 min. Quenched cells were then washed twice with DPBS and permeabilized with 0.1% Triton X-100 in PBS at room temperature for 10 min. Permeabilized cells were then washed twice with DPBS and incubated with 10 µM DBF-NHS in PBS at room temperature for 1 h, protected from light. Cells were washed 3x with PBS, then POCA was performed by incubating in 5 mM biotin-PEG3-amine (Lumiprobe Corporation 2623) in PBS at room temperature for 5 min followed by illumination with a 455-nm LED (SunLite PAR30, 15 W) at an illuminance of ∼170,000 lx at room temperature for 5 min. Cells were washed with pre-warmed (37 °C) PBST (3 x 5 min), 500 µL of 1% SDS in PBS was added to the dish, and cells were harvested by scraping into 1.5-mL microcentrifuge tubes. The biotinylated lysate was flash frozen and stored at -80 °C until further sample processing.

The lysate was thawed and crosslinks were reversed by boiling and probe sonicating according to the IF-POCA procedure. The lysate was then clarified by centrifugation at 20,000 g for 5 min, and the 500 µL supernatant was transferred to a 1.5-mL low-bind microcentrifuge tube (Fisher Scientific 02-681-320). Total protein concentration was determined by DC assay (Bio-Rad 5000112), and samples were prepared with 150 µg of protein input by diluting with PBS. For each sample, 20 μL Sera-Mag SpeedBeads Carboxyl Magnetic Beads, hydrophobic (50 μg/μL, total 1 mg) and 20 μL Sera-Mag SpeedBeads Carboxyl Magnetic Beads, hydrophilic (50 μg/μL, total 1 mg) were mixed and washed with water three times. 40 µL of 1:1 SP3 bead slurry was then transferred to each lysate and incubated at RT for 5 min with shaking (1000 RPM). Protein-bead binding was initiated by adding 800 µL of ethanol and incubating at RT for 10 min with shaking (1000 RPM), then the beads were separated from the supernatant on a magnetic rack (Sergi Lab Supplies), the supernatant discarded, and the beads washed 3x with 500 µL of 80% ethanol in UP water. The beads were then resuspended in 200 µL of freshly prepared 2 M urea and 0.5% SDS in PBS, cysteines were reduced with DTT (10 µL of a 200 mM solution, final concentration 10 mM) at 65 °C for 15 min with shaking (1000 RPM) and alkylated with iodoacetamide (10 µL of a 400 mM solution, final concentration 20 mM) at 37 °C for 30 min. Protein-bead binding was initiated by adding 600 µL of ethanol and incubating at RT for 10 min with shaking (1000 RPM), then the beads were separated from the supernatant on a magnetic rack (Sergi Lab Supplies), the supernatant discarded, and the beads washed 3x with 500 µL of 80% ethanol in UP water. The beads were resuspended in a digestion buffer consisting of 2 M urea, 4 mM calcium chloride, and 250 ng of trypsin (Worthington Biochemical LS003740) in PBS, and on-bead digest was performed at 37 °C with shaking (250 RPM) overnight.

SP3 peptide cleanup using acetonitrile and enrichment with NeutrAvidin agarose resin (50 µL/sample, Thermo Scientific 29200) were carried out as reported previously^171^. Samples were dried in a centrifugal vacuum concentrator, then reconstituted with 5% acetonitrile and 1% formic acid in UP water prior to analysis by LC-MS/MS as performed with IF-POCA.

### Fixed-Cell POCA Modified Peptide Analysis Data Processing

Raw data collected by LC–MS/MS were searched with MSFragger^166^ (v4.1) and FragPipe (v22.0) with the appropriate UniProtKB reviewed database (downloaded January 5, 2023) and same MSFragger modifications as with ISH- and IF-POCA searches. Critically, mass offsets were searched for the following modification masses: 0 (no additional modification), 388.1782 (biotin-amine), 404.1730 (biotin-sulfoxide-amine), 420.1678 (biotin-sulfone-amine).

### BioPlex Network Interaction Analysis

Proteins enriched in each condition were mapped onto the BioPlex 3.0 interaction network^16^ and interactions amongst these proteins were tallied. For this analysis, both 293T and HCT116 networks were combined to produce a single composite network. This process was then repeated against 200 randomized versions of the BioPlex 3.0 network^16^. Network randomization preserved node degree, total interactions, total vertices and bait/prey identity. Interaction counts observed across these randomized networks were used to calculate mean and standard deviations for the expected number of interactions among each protein group under the null hypothesis of interactions occurring at random, and these statistics were used to calculate z-scores for each observed interaction count. Z-scores were then used to derive p-values via a two-sided Z-test. All calculations were performed in Mathematica 14.3 (Wolfram Research Inc.).

### Statistical and Computational Analyses

#### Fourier Ring Correlation

Fourier ring correlation (FRC) analysis was performed on fluorescence microscopy images containing target signal (FISH probes) and biotinylation signal (streptavidin) for both POCA and HRP-based labeling methods. All images were reduced to two dimensions via a maximum intensity projection along the z-axis, then apodized with a two-dimensional Tukey window with a 10% cosine taper fraction to suppress edge artifacts in Fourier space. The windowed images were transformed into Fourier space via a two-dimensional discrete Fourier transform and the cross-correlation between target and biotinylation spectra was computed within concentric rings of various spatial frequencies. FRC curves as a function of feature size were constructed by averaging FRC values within each ring and smoothing with a uniform filter with a width of five frequency bins. Reported FRC curves for each labeling method show mean values across three fields of view, each containing 6-21 nuclei and 381-1273 puncta. As an upper bound determined by optical limits, an FRC curve was computed identically for an image of TetraSpeck 200nm fluorescent beads.

#### Receiver Operating Characteristic (ROC) Analysis

Receiver operating characteristic (ROC) analysis was performed using a custom script in R using the plotROC^172^ and pROC^173^ packages. True positives were proteins annotated at each subcellular location (nucleoli, nuclear speckles, nuclear envelope) in at least one of the following databases: Human Protein Atlas (HPA)^15^, CORUM^80^, and UniProtKB^81^. Proteins were ranked by their log_2_ fold change (log_2_FC) relative to a reference bait. The log_2_FC served as the continuous classifier score, with database-annotated proteins treated as the positive class and all remaining detected proteins as negatives. The area under the ROC curve (AUC) was computed for each compartment comparison, and the optimal log_2_FC threshold was identified by maximizing the Youden index (true positive rate − false positive rate). Youden values were also plotted as a function of the log_2_FC threshold to visualize classifier performance across the full score range.

#### Gene Set Enrichment Analysis (GSEA)

Gene Set Enrichment Analysis (GSEA) was performed against all Gene Ontology (GO) categories (Biological Process, Cellular Component, Molecular Function) using the gseGO function from clusterProfiler^174^, with the human annotation database org.Hs.eg.db. Proteins were ranked by log_2_FC in descending order to form the input gene list; gene sets with fewer than 10 or more than 500 members were excluded. Resulting p-values were adjusted by the Benjamini-Hochberg method. For ridge plots, the top 10 terms by adjusted p-value (ascending) across all ontologies were displayed; for bar plots, the top 5 terms were further filtered to those with NES > 0 and adjusted p-value < 0.05, ranked by −log_10_(adjusted p-value).

#### Gene Ontology Over-Representation Analysis (GO-ORA)

Gene Ontology over-representation analysis (GO-ORA) was performed using the enrichGO function from clusterProfiler^174^ across all GO ontologies (Biological Process, Cellular Component, Molecular Function). Significantly enriched proteins (log2FC > 1, Benjamini-Hochberg adjusted p-value < 0.05) were used as input and the full human annotation database served as the statistical background universe. Gene sets with fewer than 10 or more than 500 members were excluded, and p-values were adjusted using the Benjamini-Hochberg method with significance thresholds of p.adjust < 0.05 and q-value < 0.2. Bar plots display the top 5 terms ranked by ascending adjusted p-value.

#### Protein Complex Enrichment

Protein complex enrichment was performed using a custom analysis script in R using the enricherR^175^ package and CORUM^80^ database against the list of enriched proteins for each target of interest. Protein complexes were then filtered based on an adjusted p-value less than 0.01 and having at least three proteins within the enriched complex. Protein complexes that passed the designated thresholds were then visualized to depict the adjusted p value, number of enriched proteins within the enriched complex, and fraction of the total number proteins within the complex that were enriched.

## Data Availability

All supporting data for this study can be found within the article and Supplementary Information. The MS data were deposited to the ProteomeXchange Consortium with identifier PXD075788, username, and password KlT2KpznVjCv.

## Code Availability

All code used for this work is available from GitHub at github.com/SchweppeLab/poca-ms.

## Acknowledgements

We would like to thank Dr. David Shechner and members of the Backus, Beliveau, and Schweppe labs for constructive feedback and technical assistance in assembling this work. We would like to acknowledge the following sources of support: R35GM137916 (BJB), R35GM150919 (DKS), DP2GM146246 (KMB), R01AI158704 (MMHL), the W.M. Keck Foundation (BJB, DKS, KMB), Packard Fellowship (KMB), DE-FC02-02ER63421 (KMB), an Andy Hill CARE Distinguished Researcher Award (DKS), a Damon Runyon Dale Frey Award (BJB), a Cancer Consortium New Investigator Award (funded in part through P30 CA015704, DKS), the Pew Charitable Trusts (DKS), and U24HG006673 (ELH). Research reported in this publication was supported by the NHLBI under award number T32HL007093 (to CPH).

