## Supplementary Information for "A unified photosensitizer platform for *in situ* DNA-, RNA-, and protein-directed proximity labeling"

1 - Biological Chemistry Department, David Geffen School of Medicine, University of California, Los Angeles, Los Angeles, CA 90095, USA
2 - Department of Chemistry and Biochemistry, University of California, Los Angeles, Los Angeles, CA 90095, USA
3 - Department of Genome Sciences, University of Washington, Seattle, WA 98105, USA
4 - Janelia Research Campus, Howard Hughes Medical Institute, Ashburn, VA 20147, USA
5 - Department of Cell Biology, Harvard Medical School, Boston, MA 02115, USA
6 - Brotman Baty Institute for Precision Medicine, Seattle, WA 98105, USA
7 - Institute for Stem Cell & Regenerative Medicine, University of Washington, Seattle, WA 98105, USA
8 - Molecular Biology Institute, University of California, Los Angeles, Los Angeles, CA 90095, USA
9 - UCLA-DOE Institute for Genomics and Proteomics, University of California, Los Angeles, Los Angeles, CA 90095, USA
10 - Jonsson Comprehensive Cancer Center, University of California, Los Angeles, Los Angeles, CA 90095, USA
11 - Eli and Edythe Broad Center of Regenerative Medicine and Stem Cell Research, University of California, Los Angeles, Los Angeles, CA 90095, USA
12 - These authors contributed equally: Elijah B Biletch, Conor P Herlihy, and Lidan Li

### Table of Contents

**1. Supplementary Figures (pp. 3–20)**
**Figure S1**. DBF-NHS and $JF_{570}$-NHS synthesis and conjugation to oligonucleotide and antibody probes.
**Figure S2**. Validation of POCA labeling conditions.
**Figure S3**. POCA labeling across diverse cell lines spanning species and culture formats.
**Figure S4**. Labeled peptide analysis confirms a histidine-based biotin-amine POCA labeling mechanism.
**Figure S5**. ISH-POCA reveals nucleotide specific protein interactions.
**Figure S6**. IF-POCA reveals NPM1, SON, and NUP98 proximal proteomes.
**Figure S7**. POCA labeling is adaptable to different photosensitizers.
**Figure S8**. IF-POCA proteomics captures the interactome of additional endogenous protein targets.
**Figure S9**. Pairing ISH- and IF-POCA provides high-confidence interactome maps.

**2. Supplementary Tables (pp. 21–23)**
**Table S1**. Reagents used for ISH- and IF-POCA.
**Table S2**. Primary antibodies used for IF-POCA.
**Table S3**. List of proteomics raw files used in this report.
**Table S4**. Sequences of oligonucleotides used for ISH-POCA.
**Table S5**. Raw and processed mass spectrometry data tables corresponding to Figure 2
**Table S6**. Raw and processed mass spectrometry data tables corresponding to Figure 3
**Table S7**. Raw and processed mass spectrometry data tables corresponding to Figure S8

**3. Supplementary Note: Synthetic schemes and procedures, NMRs (pp. 24–30)**

**4. References (p. 31)**

### 1. Supplementary Figures

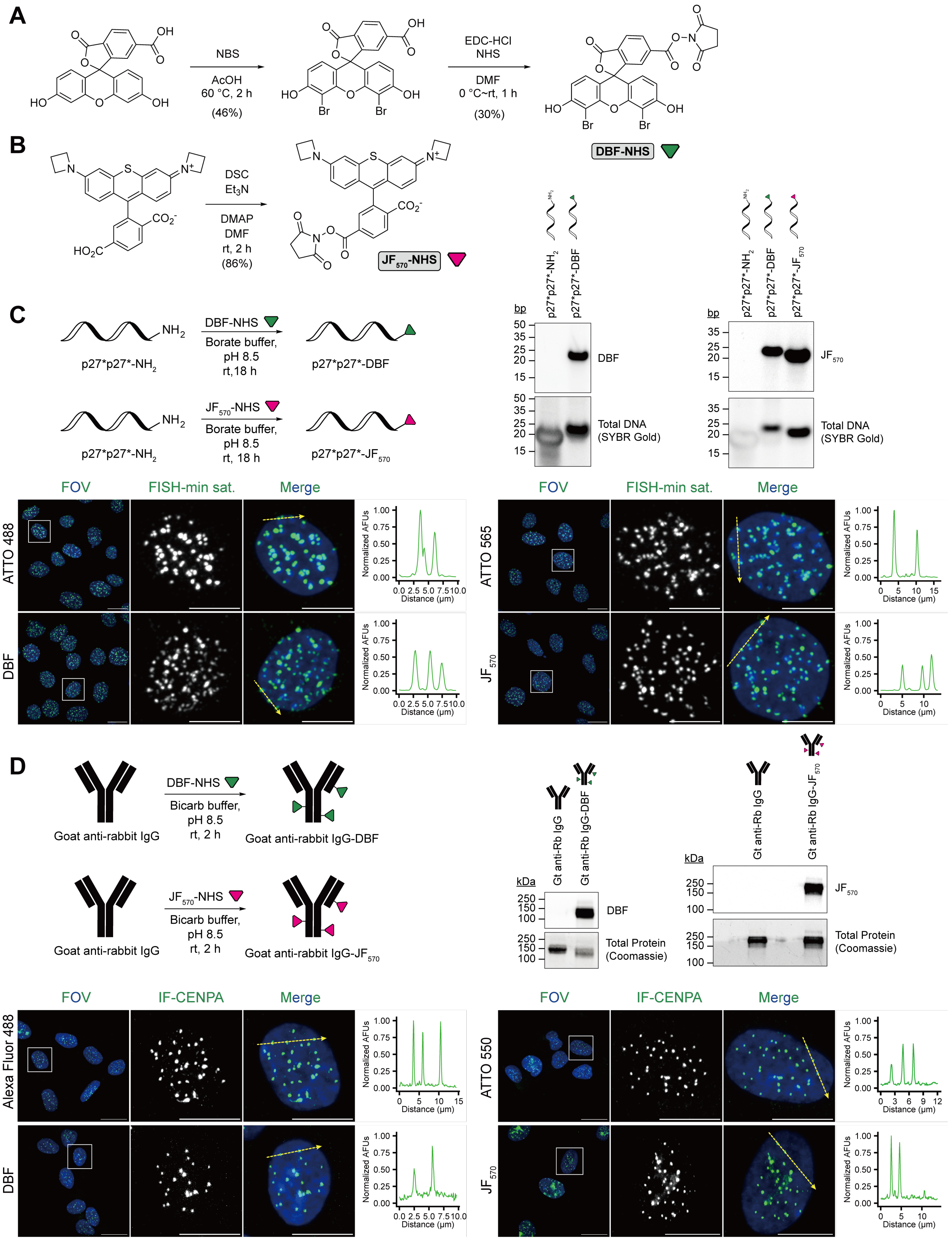

**Figure S1. DBF-NHS and $JF_{570}$-NHS synthesis and conjugation to oligonucleotide and antibody probes. A)** Synthesis of 6-carboxy-DBF NHS ester (DBF-NHS) was achieved in two steps by bromination of 6-carboxyfluorescein (6-FAM) followed by EDC-mediated coupling of *N*-hydroxysuccinimide. **B)** Synthesis of 6-carboxy-$JF_{570}$ NHS ester ($JF_{570}$-NHS) was achieved by DSC-mediated esterification of 6-carboxy-$JF_{570}$. **C)** Conjugation of an amine-modified oligonucleotide ISH probe with DBF-NHS or $JF_{570}$-NHS assayed by native PAGE gel via in-gel DBF or $JF_{570}$ fluorescence and SYBR Gold staining and FISH validation. DBF signal is observed in the $JF_{570}$ in-gel fluorescence channel due to fluorophore spectral overlap. Representative images are of a merged image with DAPI (blue), minor satellite FISH (green), and a cropped image of a single cell from the merged image. p27*p27* refers to the following oligonucleotide sequence: 5'-Amine-$C_6$-TTTATGATGATGTATGATGATGT-3'[1]. **D)** Conjugation of a secondary antibody IF probe with DBF-NHS or $JF_{570}$-NHS assayed by non-reducing SDS-PAGE gel via in-gel DBF or $JF_{570}$ fluorescence and Coomassie blue staining and IF validation. Representative images of a merged image with DAPI (blue), CENPA IF (green), and a cropped image of a single cell from the merged image. For **C** & **D**, all images are maximum intensity projections. Scale bars are 20 μm in the standard images and 10 μm in the cropped images. Depiction of the line scan through foci are shown to the right of cropped images. Line scan graphs show fluorescence intensity of FISH/IF along the lines in the adjacent representative images.

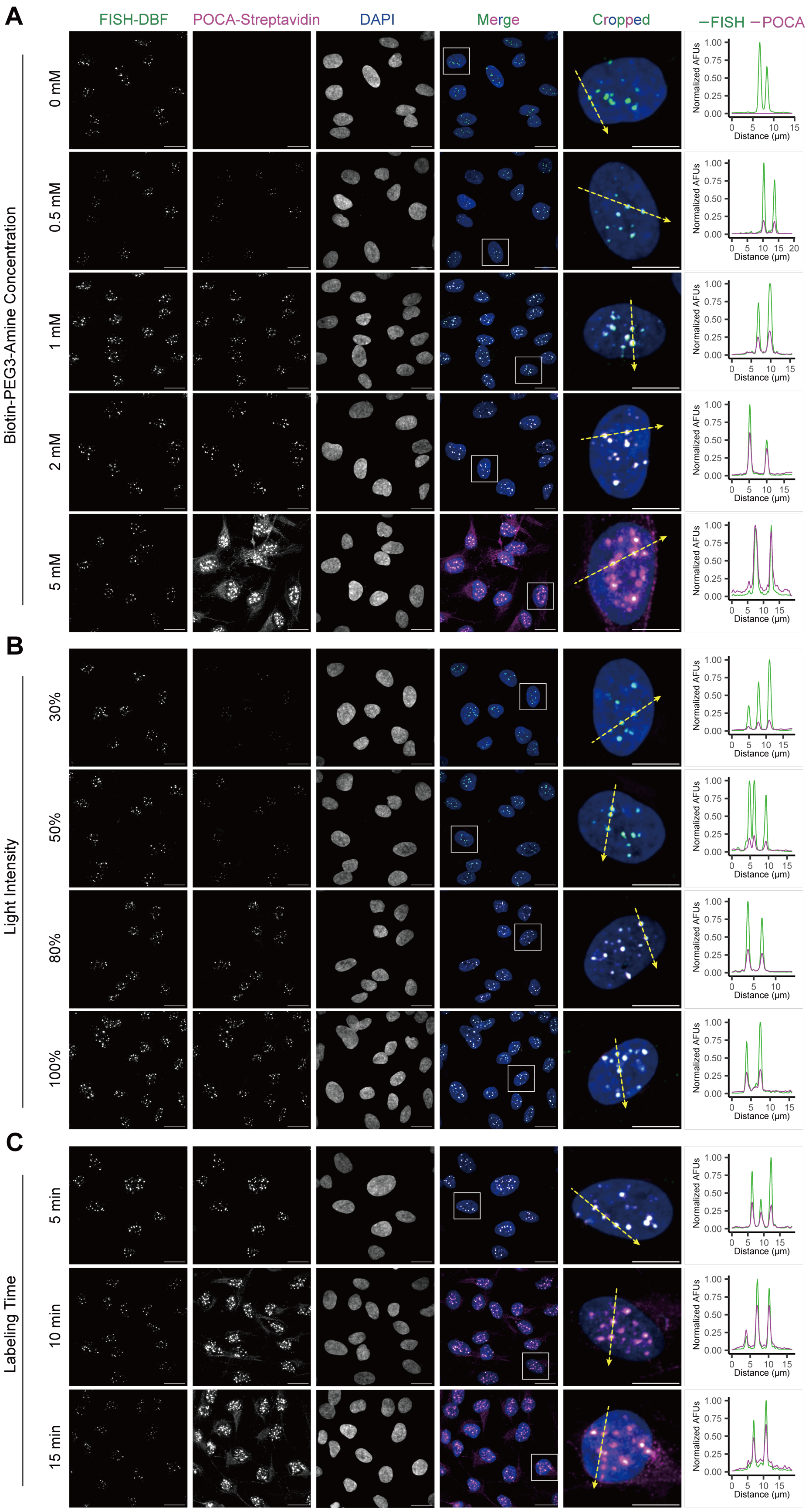

A
B
C
FISH-DBF
POCA-Streptavidin
DAPI
Merge
Cropped
−FISH −POCA
Biotin-PEG3-Amine Concentration
0 mM
0.5 mM
1 mM
2 mM
5 mM
Light Intensity
30%
50%
80%
100%
Labeling Time
5 min
10 min
15 min
Normalized AFUs
Distance (μm)

**Figure S2. Validation of POCA labeling conditions. A, B, C)** Representative images of Pan Alpha ISH-POCA labeling depicting FISH-DBF (green), POCA-Streptavidin (magenta) signal, a merged image of both labeling channels with DAPI (blue), and a cropped image of a single cell from the merged image. Overlap between DBF and streptavidin signal appears white. All images are maximum intensity projections. Scale bars are 20 μm in the standard images and 10 μm in the cropped images. Depiction of the line scan through Pan Alpha foci are shown to the right of cropped images. Line scan graphs show fluorescence intensity of DBF (green) and streptavidin (magenta) along the lines in the adjacent representative images. **A** Shows altered concentrations of Biotin used for labeling, **B** shows variation in light intensity during the labeling reaction, and **C** shows variable labeling time.

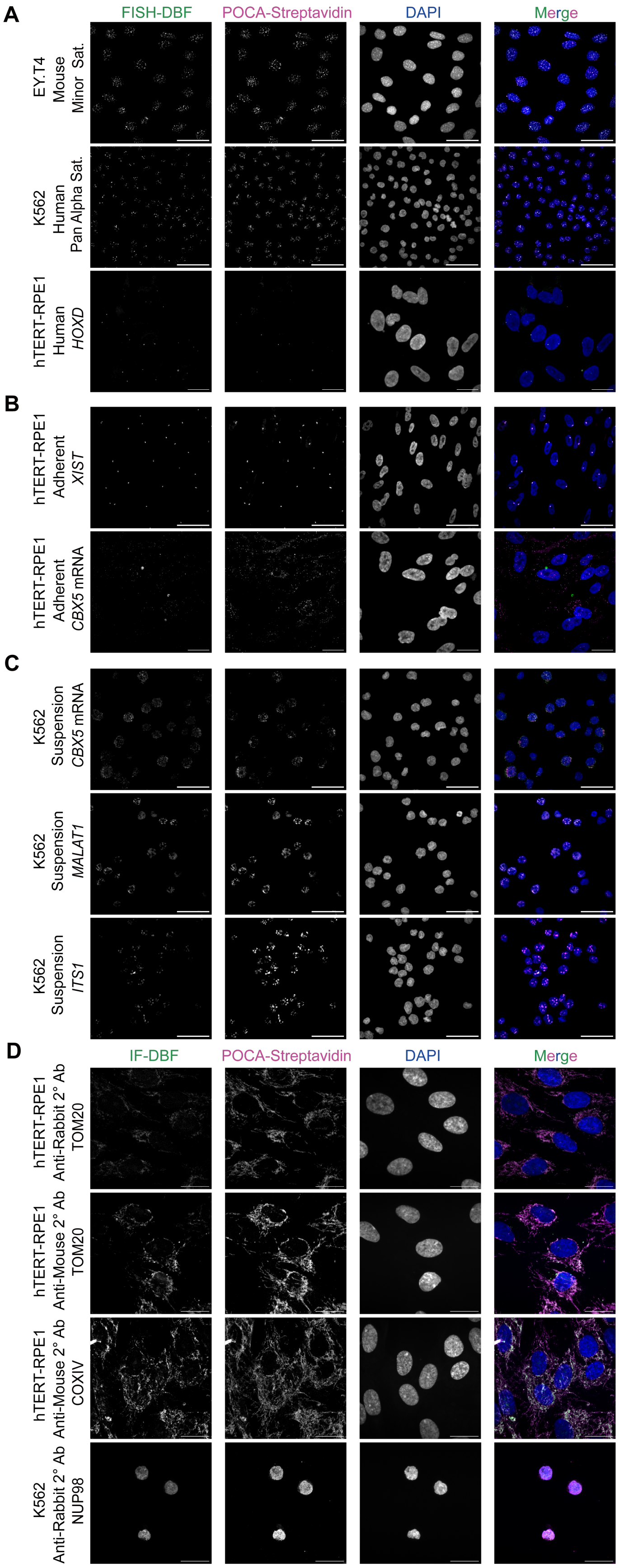
A
FISH-DBF
POCA-Streptavidin
DAPI
Merge
EY.T4
Mouse
Minor Sat.
K562
Human
Pan Alpha Sat.
hTERT-RPE1
Human
HOXD
B
hTERT-RPE1
Adherent
XIST
hTERT-RPE1
Adherent
CBX5 mRNA
C
K562
Suspension
CBX5 mRNA
K562
Suspension
MALAT1
K562
Suspension
ITS1
D
IF-DBF
POCA-Streptavidin
DAPI
Merge
hTERT-RPE1
Anti-Rabbit 2° Ab
TOM20
hTERT-RPE1
Anti-Mouse 2° Ab
TOM20
hTERT-RPE1
Anti-Mouse 2° Ab
COXIV
K562
Anti-Rabbit 2° Ab
NUP98

**Figure S3. POCA labeling across diverse cell lines spanning species and culture formats. A)** Representative images of ISH-POCA labeling of minor satellite repeats in EY.T4 mouse embryonic fibroblast (EY.T4) cells (60x), alpha satellite repeats in human myelogenous leukemia (K562) suspension cells (60x), and *HOXD* genomic DNA in human retinal pigment epithelial (hTERT-RPE1) cells (100x). **B)** Representative images of ISH-POCA labeling of *XIST* (60x) and *CBX5* mRNA (100x) in hTERT-RPE1 cells. **C)** Representative images taken at 60x of ISH-POCA labeling of *CBX5*, *MALAT1* and *ITS1* in K562 suspension cells. **D)** Representative images taken at 100x of IF-POCA labeling using the DBF-conjugated anti-rabbit/mouse IgG secondaries of TOM20 and COXIV in hTERT-RPE1 cells, and NUP98 in K562 cells. All images are maximum intensity projections with FISH/IF-DBF (green), POCA-Streptavidin (magenta), and a merged image of both labeling channels with DAPI (blue). Overlap between DBF and streptavidin signal appears white. Scale bars are 50 μm in the 60x images and 20 μm in the 100x images.

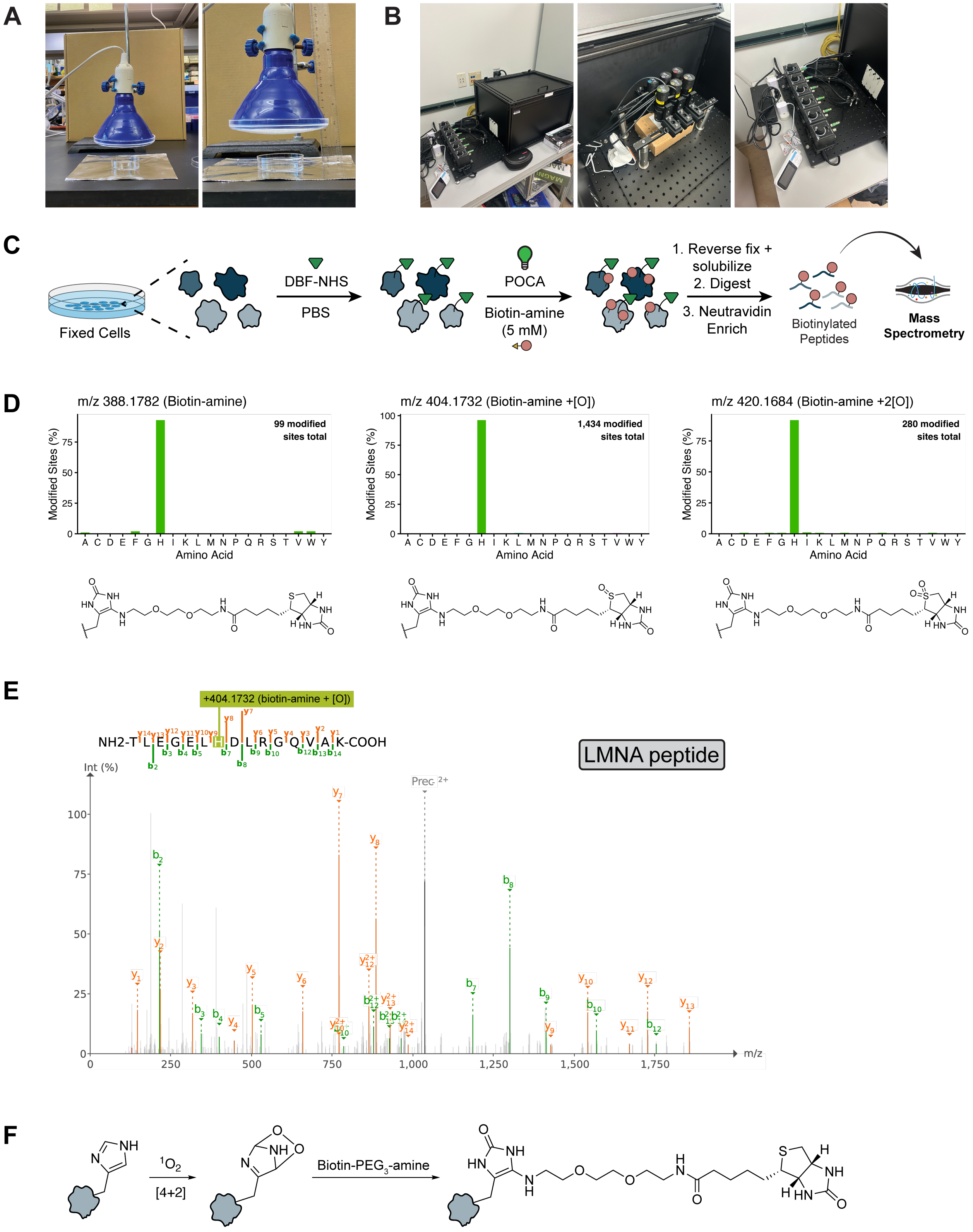


**Figure S4. Labeled peptide analysis confirms a histidine-based biotin-amine POCA labeling mechanism. A & B)** Pictures showing the UCLA (A) and UW (B) POCA labeling illumination platforms used for POCA proteomics workflows. **C)** Schematic showing the workflow for fixed whole-cell POCA biotinylated peptide analysis. **D)** Biotin-amine

POCA amino acid labeling preference via FragPipe mass offset search using modification masses for biotin-amine, singly oxidized biotin-amine (biotin-sulfoxide-amine), and doubly oxidized biotin-amine (biotin-sulfone-amine). The bar graph represents the fraction of all modified sites that were modified at the indicated amino acid. **E)** Annotated LMNA peptide MS2 spectrum showing the assigned localization site using a closed search with the respective fixed modification at histidine. **F)** Proposed mechanism of histidine oxidation and capture by biotin-$PEG_3$-amine.

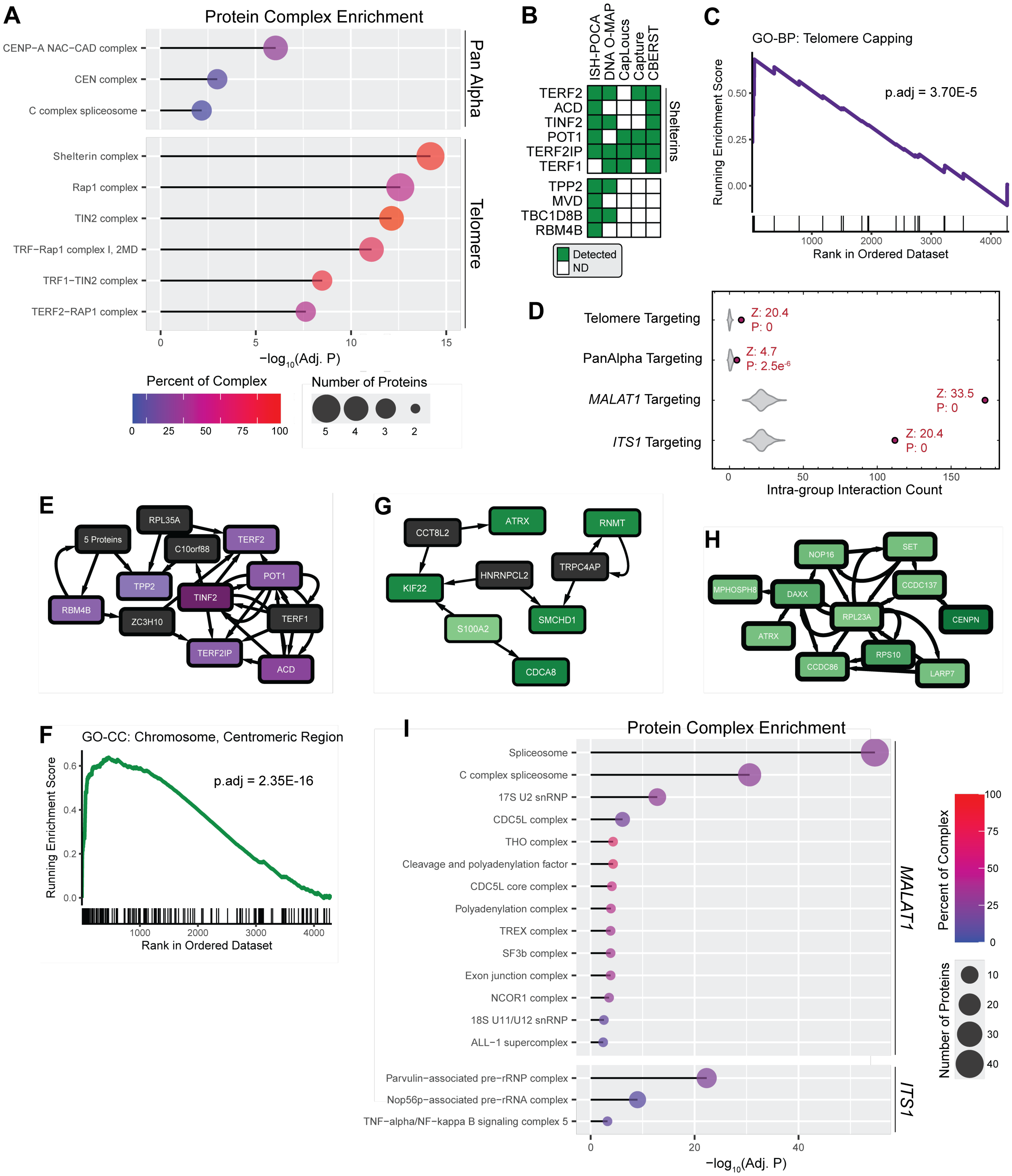

**Figure S5. ISH-POCA reveals nucleotide-specific protein interactions. A)** Protein complex enrichment analysis using the CORUM[2] database of Pan Alpha and Telomere-enriched proteins. Dot size is a function of the total number of proteins enriched from the complex, dot color is the percent of proteins from the complex that were enriched, and line length is a function of the -$Log_{10}$(adjusted p-value). Adjusted p-value < 0.05. **B)** Heatmap of proteins identified by Telomere ISH-POCA and previous studies. **C)** Gene Set Enrichment Analysis (GSEA) for Gene Ontology (GO) term "telomere capping" (GO:0016233) in the Pan Alpha vs Telomere ISH-POCA comparison. The colored lines show the running enrichment score (ES) traversing the ranked list of proteins by fold-change. Black tick marks indicate the positions of member proteins within the ranked list. The peak ES corresponds to the normalized enrichment score (NES). **D)** Analysis of protein-protein interactions beyond individual complex membership based on the BioPlex[3,4] interactome networks of Pan Alpha, Telomere, *MALAT1,* and *ITS1*, respectively. **E)** BioPlex interaction network of significantly enriched proteins at Telomeres. Connections represent detected protein-protein interactions by AP-MS with color represented as a function of $\log_2$ fold-change by ISH-POCA with black boxes indicating proteins that were not detected. **F)** Gene Set Enrichment Analysis (GSEA) for Gene Ontology (GO) term "chromosome, centromeric region" (GO:0000775) in the Pan Alpha vs Telomere ISH-POCA comparison. **G, H)** BioPlex interaction networks of significantly enriched proteins at Pan Alpha. Connections represent detected protein-protein interactions by AP-MS with color represented as a function of $\log_2$ fold-change by ISH-POCA with black boxes indicating proteins that were not detected. **I)** Protein complex enrichment analysis using the CORUM[2] database of *MALAT1* and *ITS1* enriched proteins. Dot size is a function of the total number of proteins enriched from the complex, dot color is the percent of proteins from the complex that were enriched, and line length is a function of the $-\text{Log}_{10}$(adjusted p-value). Adjusted p-value $< 0.01$.

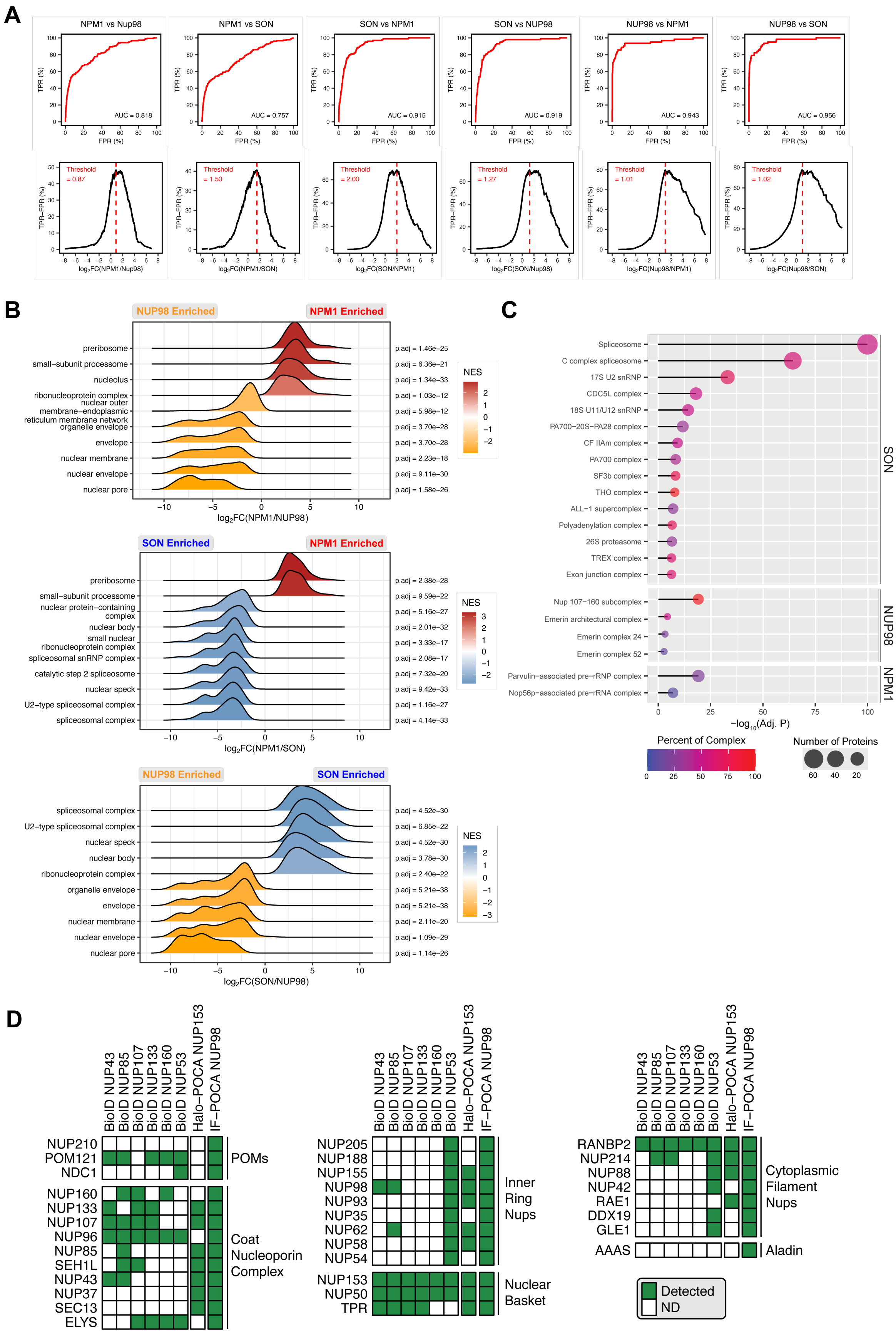

A
NPM1 vs Nup98
NPM1 vs SON
SON vs NPM1
SON vs NUP98
NUP98 vs NPM1
NUP98 vs SON
AUC = 0.818
AUC = 0.757
AUC = 0.915
AUC = 0.919
AUC = 0.943
AUC = 0.956
TPR (%)
FPR (%)
TPR−FPR (%)
Threshold = 0.87
Threshold = 1.50
Threshold = 2.00
Threshold = 1.27
Threshold = 1.01
Threshold = 1.02
B
NUP98 Enriched
NPM1 Enriched
SON Enriched
NES
log2FC(NPM1/NUP98)
log2FC(NPM1/SON)
log2FC(SON/NUP98)
C
Percent of Complex
Number of Proteins
−log10(Adj. P)
D
Detected
ND
POMs
Coat Nucleoporin Complex
Inner Ring Nups
Nuclear Basket
Cytoplasmic Filament Nups
Aladin

**Figure S6. IF-POCA reveals NPM1, SON, and NUP98 proximal proteomes. A)** Receiver operating characteristic (ROC) curves for each IF-POCA pairwise bait comparison. Proteins were ranked in descending order according to fold-change, and each ranked protein was used as a classification threshold to compute true positive rate (TPR) and false positive rate (FPR) based on subcellular annotations in at least two of the following: the Human Protein Atlas (HPA)[5], CORUM[2], and UniProtKB[6]. **B)** Ridge plots showing the top 10 enriched Gene Ontology Cellular Component (GO-CC) terms, ranked by Benjamini-Hochberg adjusted p-value, from Gene Set Enrichment Analysis (GSEA) for IF-POCA pairwise comparisons between the three baits. Each ridge represents the density distribution of ranked genes within a given GO-CC term, with color indicating the normalized enrichment score (NES). For GSEA, proteins were ranked according to fold-change. **C)** Protein complex enrichment analysis using the CORUM[2] database of SON, NUP98, and NPM1 enriched proteins. Dot size is a function of the total number of proteins enriched from the complex (Number of Proteins), dot color is the percent of proteins from the complex that were enriched (Percent of Complex), and line length is a function of the $-\text{Log}_{10}$(adjusted P-value). Adjusted P-value < 0.05. **D)** Heatmap of proteins identified by NUP98 IF-POCA and previous nuclear pore proximity labeling studies using BioID[7] and Halo-POCA[8]. "Detected" indicates at least one spectral count.

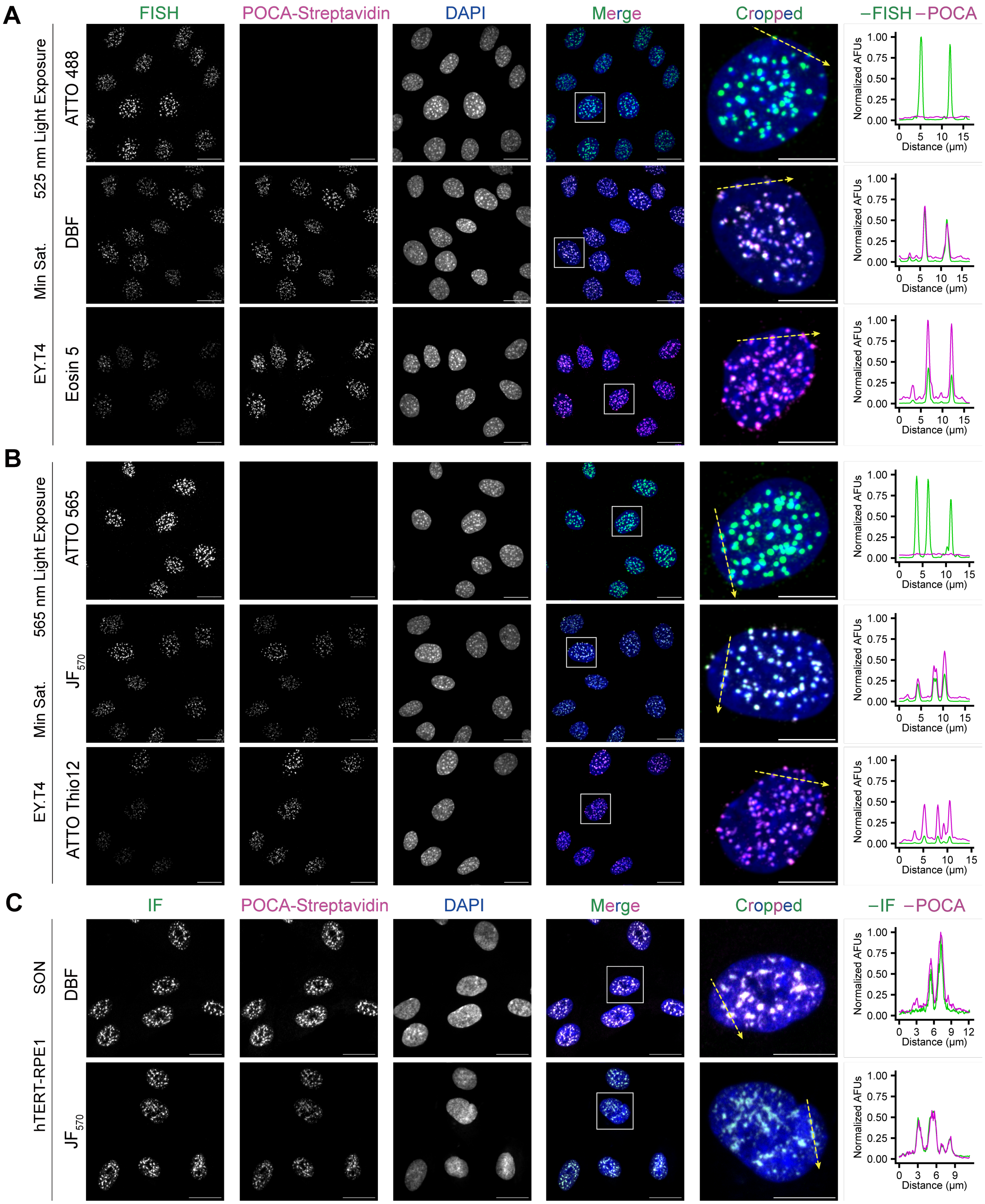


**Figure S7. POCA labeling is adaptable to different photosensitizers. A,B)** ISH-POCA labeling of minor satellite loci in EY.T4 cells. **A** 525-nm irradiation: ATTO 488 shows no detectable POCA labeling, whereas DBF and Eosin-5 produce clear labeling. **B** 565-nm irradiation: ATTO 565 shows no detectable labeling, whereas $JF_{570}$ and ATTO Thio 12 produce detectable POCA labeling. **C)** IF-POCA labeling of SON in hTERT-RPE1 cells using DBF and JF570. Representative maximum-intensity projections show FISH or IF (green), POCA-streptavidin (magenta), merged images with DAPI (blue), and cropped single-cell views; colocalization appears white. Line scans across representative foci show fluorescence intensity profiles of FISH (green) and streptavidin (magenta) along the dotted yellow lines. Scale bars, 20 μm (full images) and 10 μm (cropped images).

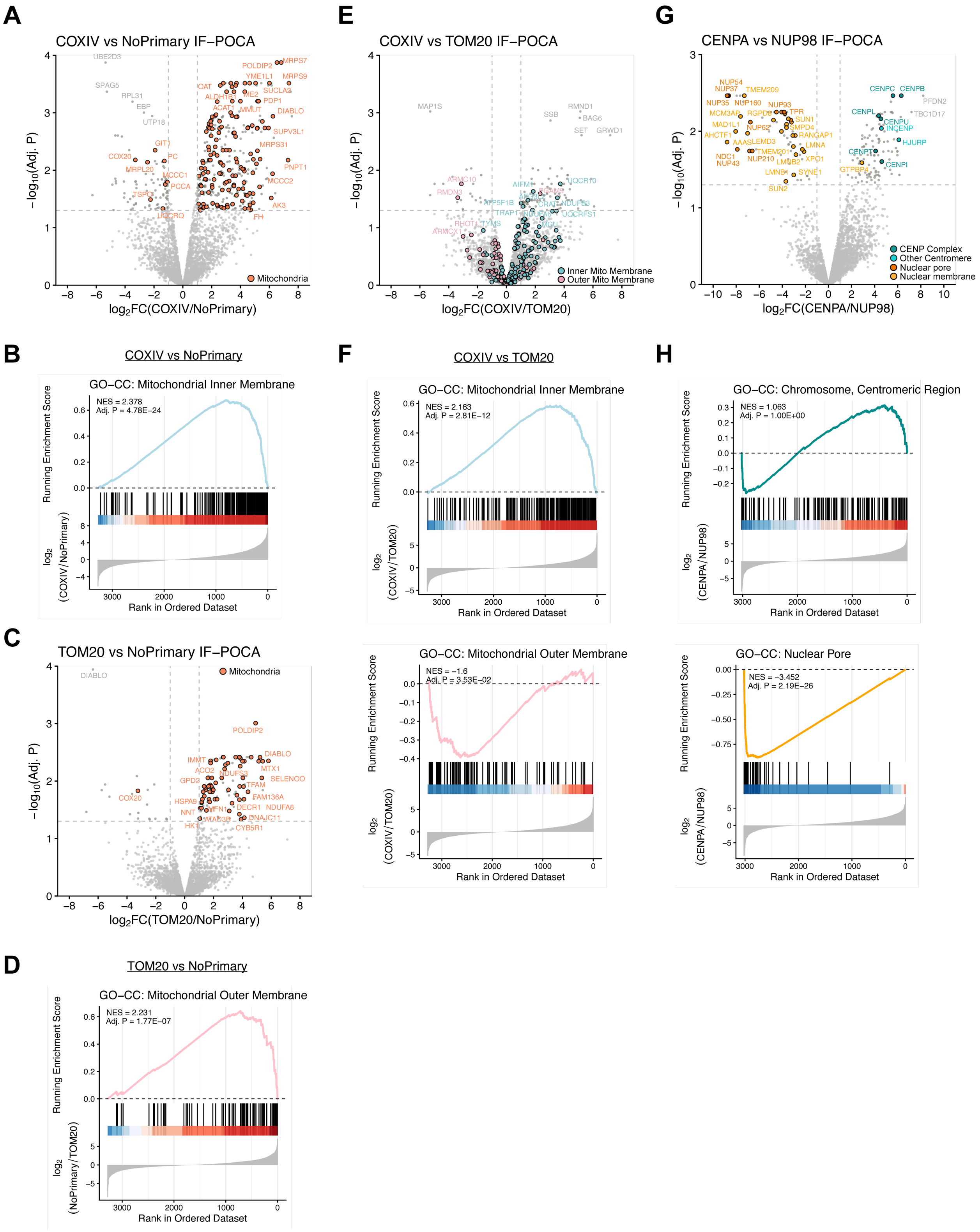


**Figure S8. IF-POCA proteomics captures the interactome of additional endogenous protein targets. A)** Volcano plot of proteins identified in IF-POCA proteomics pairwise comparison targeting COXIV or omitting primary antibody. Each dot represents a single protein. Mitochondrial proteins are labeled salmon based on UniProtKB subcellular annotation[6]. Hashed lines delineate the thresholds for significance, with greater than 2-fold change in relative enrichment and a Benjamini-Hochberg adjusted P value less than 0.05. **B)** Gene Set Enrichment Analysis (GSEA) running enrichment score plot for the Gene Ontology Cellular Component (GO-CC) term “mitochondrial inner membrane” (GO:0005743) in the COXIV vs NoPrimary experiment. The colored lines show the running enrichment score (ES) traversing the ranked list of proteins by fold-change. Black tick marks indicate the positions of member proteins within the ranked list. The peak ES corresponds to the normalized enrichment score (NES). The red-to-blue color gradient represents the $\log_2$ fold-change ranking metric, where red denotes higher $\log_2$ fold-change and blue denotes lower $\log_2$ fold-change. **C)** Volcano plot of proteins identified in IF-POCA proteomics pairwise comparison targeting TOM20 or omitting primary antibody. **D)** GSEA running enrichment score plot for the GO-CC term “mitochondrial outer membrane” (GO:0005741) in the TOM20 vs NoPrimary experiment. **E)** Volcano plot of proteins identified in IF-POCA proteomics pairwise comparison targeting COXIV or TOM20. Proteins are labeled light blue for inner mitochondrial membrane and pink for outer mitochondrial membrane based on UniProtKB subcellular annotation. **F)** GSEA running enrichment score plots for the GO-CC terms “mitochondrial inner membrane” (GO:0005743) and “mitochondrial outer membrane” (GO:0005741) in the COXIV vs TOM20 comparison. **G)** Volcano plot of proteins identified in IF-POCA proteomics pairwise comparison targeting CENPA or NUP98. Proteins of interest are labeled cyan for centromeres, orange for nuclear envelope, and brown for nuclear pore. **H)** GSEA running enrichment score plots for the GO-CC terms “chromosome, centromeric region” (GO:0000775) and “nuclear pore” (GO:0005643) in the CENPA vs NUP98 comparison. All raw and processed proteomics data associated with this figure can be found in Table S7.

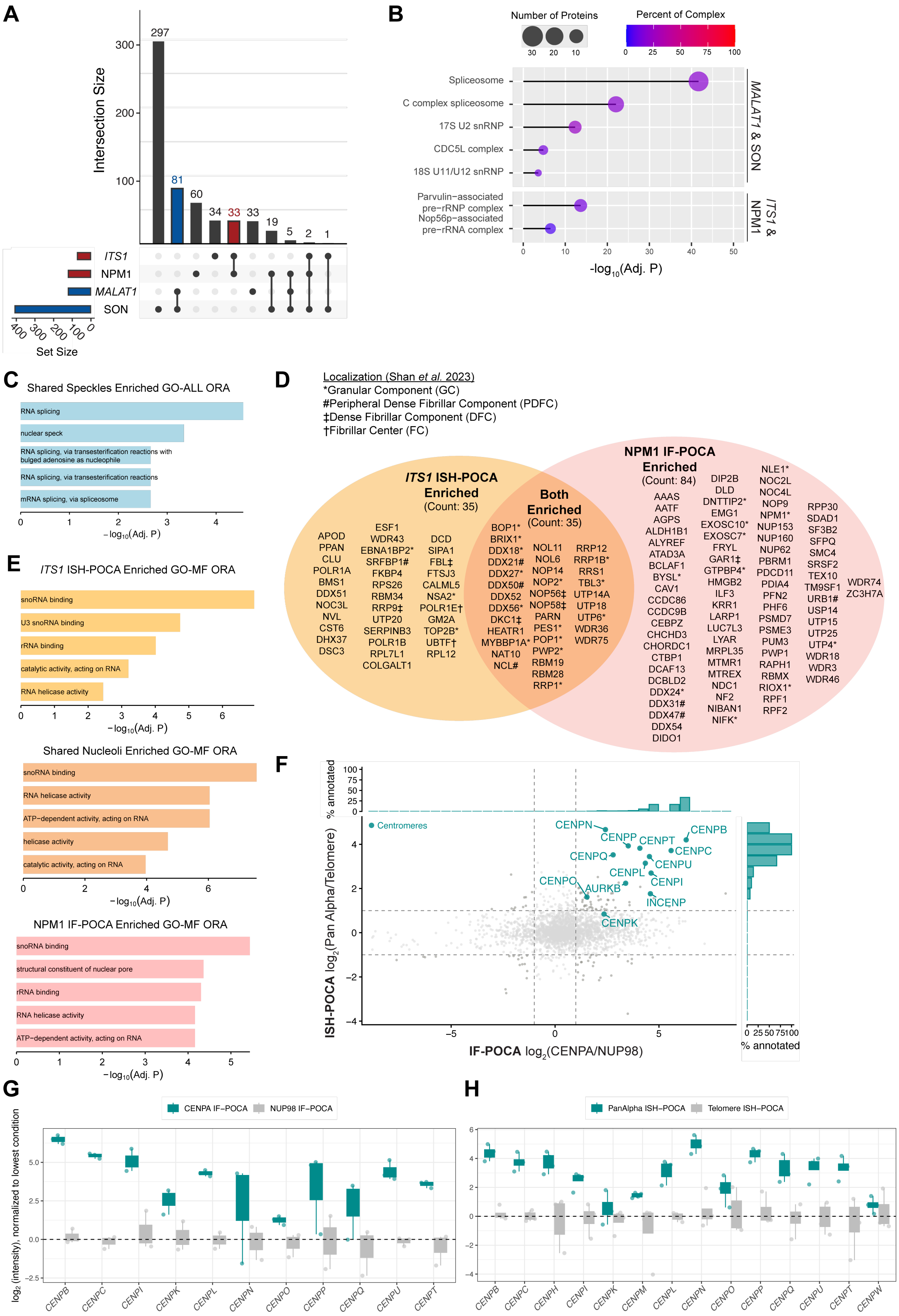
A
Intersection Size
297
81
60
34
33
33
19
5
2
1
ITS1
NPM1
MALAT1
SON
Set Size
B
Number of Proteins
Percent of Complex
Spliceosome
C complex spliceosome
17S U2 snRNP
CDC5L complex
18S U11/U12 snRNP
Parvulin−associated pre−rRNP complex
Nop56p−associated pre−rRNA complex
MALAT1 & SON
ITS1 & NPM1
-log10(Adj. P)
C
Shared Speckles Enriched GO-ALL ORA
RNA splicing
nuclear speck
RNA splicing, via transesterification reactions with bulged adenosine as nucleophile
RNA splicing, via transesterification reactions
mRNA splicing, via spliceosome
−log10(Adj. P)
D
Localization (Shan et al. 2023)
*Granular Component (GC)
#Peripheral Dense Fibrillar Component (PDFC)
‡Dense Fibrillar Component (DFC)
†Fibrillar Center (FC)
ITS1 ISH-POCA Enriched (Count: 35)
APOD PPAN CLU POLR1A BMS1 DDX51 NOC3L NVL CST6 DHX37 DSC3
ESF1 WDR43 EBNA1BP2* SRFBP1# FKBP4 RPS26 RBM34 RRP9‡ UTP20 SERPINB3 POLR1B RPL7L1 COLGALT1
DCD SIPA1 FBL‡ FTSJ3 CALML5 NSA2* POLR1E† GM2A TOP2B* UBTF† RPL12
Both Enriched (Count: 35)
BOP1* BRIX1* DDX18* DDX21# DDX27* DDX50# DDX52 DDX56* DKC1‡ HEATR1 MYBBP1A* NAT10 NCL#
NOL11 NOL6 NOP14 NOP2* NOP56‡ NOP58‡ PARN PES1* POP1* PWP2* RBM19 RBM28 RRP1*
RRP12 RRP1B* RRS1 TBL3* UTP14A UTP18 UTP6* WDR36 WDR75
NPM1 IF-POCA Enriched (Count: 84)
AAAS AATF AGPS ALDH1B1 ALYREF ATAD3A BCLAF1 BYSL* CAV1 CCDC86 CCDC9B CEBPZ CHCHD3 CHORDC1 CTBP1 DCAF13 DCBLD2 DDX24* DDX31# DDX47# DDX54 DIDO1
DIP2B DLD DNTTIP2* EMG1 EXOSC10* EXOSC7* FRYL GAR1‡ GTPBP4* HMGB2 ILF3 KRR1 LARP1 LUC7L3 LYAR MRPL35 MTMR1 MTREX NDC1 NF2 NIBAN1 NIFK*
NLE1* NOC2L NOC4L NOP9 NPM1* NUP153 NUP160 NUP62 PBRM1 PDCD11 PDIA4 PFN2 PHF6 PSMD7 PSME3 PUM3 PWP1 RAPH1 RBMX RIOX1* RPF1 RPF2
RPP30 SDAD1 SF3B2 SFPQ SMC4 SRSF2 TEX10 TM9SF1 URB1# USP14 UTP15 UTP25 UTP4* WDR18 WDR3 WDR46
WDR74 ZC3H7A
E
ITS1 ISH-POCA Enriched GO-MF ORA
snoRNA binding
U3 snoRNA binding
rRNA binding
catalytic activity, acting on RNA
RNA helicase activity
−log10(Adj. P)
Shared Nucleoli Enriched GO-MF ORA
snoRNA binding
RNA helicase activity
ATP−dependent activity, acting on RNA
helicase activity
catalytic activity, acting on RNA
−log10(Adj. P)
NPM1 IF-POCA Enriched GO-MF ORA
snoRNA binding
structural constituent of nuclear pore
rRNA binding
RNA helicase activity
ATP−dependent activity, acting on RNA
−log10(Adj. P)
F
% annotated
Centromeres
CENPN
CENPP
CENPT
CENPB
CENPC
CENPQ
CENPU
CENPL
CENPO
CENPI
AURKB
INCENP
CENPK
ISH-POCA log2(Pan Alpha/Telomere)
IF-POCA log2(CENPA/NUP98)
% annotated
G
CENPA IF−POCA
NUP98 IF−POCA
log2 (intensity), normalized to lowest condition
CENPB CENPC CENPI CENPK CENPL CENPN CENPO CENPP CENPQ CENPU CENPT
H
PanAlpha ISH−POCA
Telomere ISH−POCA
CENPB CENPC CENPH CENPI CENPK CENPM CENPL CENPN CENPO CENPP CENPQ CENPU CENPT CENPW

**Figure S9. Pairing ISH- and IF-POCA provides high-confidence interactome maps. A)** Upset plot depicting the number of shared and uniquely enriched proteins across all four baits used for IF-POCA and ISH-POCA targeting of nucleoli and nuclear speckles. **B)** Protein complex enrichment analysis using the CORUM[2] database of enriched proteins shared between *MALAT1* and SON or *ITS1* and NPM1. Dot size is a function of the total number of proteins enriched from the complex, dot color is the percent of proteins from the complex that were enriched, and line length is a function of the $-\log_{10}$(adjusted p-value). Adjusted p-value < 0.05. **C)** Gene Ontology with Molecular Function, Biological Process, and Cellular Component terms (GO-ALL) over-representation analysis (ORA) of proteins significantly enriched ($\log_2$FC > 1, Benjamini-Hochberg adjusted P-value < 0.05) in both *MALAT1* ISH-POCA and SON IF-POCA versus the remainder of the human proteome. The x-axis corresponds to the $-\log_{10}$ Benjamini-Hochberg adjusted P-value for ORA of each term. **D)** Complete lists of proteins significantly enriched ($\log_2$FC > 1, Benjamini-Hochberg adjusted P-value < 0.05) in uniquely *ITS1* ISH-POCA (versus *MALAT1*) (orange), in uniquely NPM1 IF-POCA (versus SON) (red), and in both analyses (red-orange). Asterisk annotations correspond to proteins with super-resolution microscopy validated localizations in the respective sub-nucleolar layers, via Shan et al. 2023[9]. **E)** Gene Ontology Molecular Function (GO-MF) ORA of each protein list versus the remainder of the human proteome. The x-axis corresponds to the $-\log_{10}$ Benjamini-Hochberg adjusted P-value for ORA of each term. **F)** Enrichment distribution of known nucleolar and nuclear speckle proteins as defined by at least one of the HPA[5], CORUM[2], and UniProtKB[6] databases. Dashed lines denote the threshold of enrichment, $\log_2$ fold-change > 2. Histograms depict the fraction of proteins in a particular range of $\log_2$ fold-change of width 0.5 that have annotations for centromeres (cyan) for each method, expressed as a percentage. **G,H)** Box-and-whisker plots showing relative $\log_2$ protein intensities of CENP family proteins in CENPA vs NUP98 IF-POCA (**G**) or PanAlpha vs Telomere ISH-POCA (**H**). For each protein, $\log_2$ intensities are normalized by subtracting the median $\log_2$ intensity of the lowest signal condition, such that the y-axis reflects $\log_2$ fold-enrichment relative to the per-protein baseline. Boxes span the interquartile (IQL) range, whiskers extend to 1.5 x IQL.

### 2. Supplementary Tables

**Table S1.** Reagents used for ISH- and IF-POCA.

| Reagent | Source | Identifier | Cost (USD) |
|---|---|---|---|
| **Chemicals** | | | |
| Biotin-PEG3-amine | Lumiprobe | 2623-100 mg | $119 / 100 mg |
| Propargylamine | Combi-Blocks | OS-7456 | $10 / 5 g |
| DBF-NHS | *In-house prepared (Figure S1A)* | | ~$200–300 / 1 g |
| $JF_{570}$-NHS | *In-house prepared (Figure S1B)* | | |
| **Oligonucleotides** | | | |
| p27*p27*-C6-$NH_2$ | IDT | N/A | $38 / 25 nmol |
| p27*p27*-C6-DBF | *In-house prepared (Figure S1C)* | | ~$25 / 10 nmol |
| p27*p27*-ATTO 565 | IDT | N/A | $250 / 100 nmol |
| p27*p27*-ATTO 488 | IDT | N/A | $280 / 100 nmol |
| p27*p27*-ATTO Thio12 | BioSynthesis | N/A | $995 / 5 nmol |
| p27*p27*-Eosin 5 | BioSynthesis | N/A | $595 / 5 nmol |
| **Antibodies** | | | |
| Goat anti-rabbit IgG (H+L) | Thermo Fisher | A16098 | $299 / 2 mg |
| Goat anti-rabbit IgG (H+L), DBF-conjugated | *In-house prepared (Figure S1D)* | | ~$75 / 300 µg |
| Goat anti-mouse IgG (H+L) | Abcam | ab6708 | $175 / 1 mg |
| Goat anti-mouse IgG (H+L), DBF-conjugated | *In-house prepared (Figure S1D)* | | ~$60 / 300 µg |
| **Irradiation Setup** | | | |
| Sunlite PAR30 blue LED (8 watt) - UCLA Platform | Sunlite | 81472-SU | $8 per lamp |
| 508 nm, 550 mW (Min) Mounted LED, 1200 mA - UW Platform | Thor Labs | M505L5 | $275.63 per lamp |
| **Estimated cost per sample replicate (~2 million cells)** | | | |
| **ISH-POCA** | **Total cost ~ $151.14 per sample replicate** | | |
| | ~$1.14 for POCA reagents (p27*p27*-C6-DBF + biotin-PEG3-amine) | | |
| | ~$50 for proteomics sample preparation | | |
| | ~$100 for proteomics instrument time per sample replicate | | |
| **IF-POCA** | **Total cost ~ $152.60 per sample replicate** | | |
| | ~$2.60 for POCA reagents (DBF-conjugated anti-Rb IgG + biotin-PEG3-amine) | | |
| | ~$50 for proteomics sample preparation | | |
| | ~$100 for proteomics instrument time per sample replicate | | |

**Table S2.** Primary antibodies used for IF-POCA.

| Reagent | Source | Identifier | Cost (USD) |
|---|---|---|---|
| **Primary Antibodies** | | | |
| Rabbit anti-NPM1 | Cell Signaling Technology | 92825 | $415 / 100 μL |
| Rabbit anti-SON | Atlas Antibodies | HPA023535 | $530 / 100 μL |
| Rabbit anti-NUP98 | Cell Signaling Technology | 2598 | $339 / 100 μL |
| Rabbit anti-CENPA | Cell Signaling Technology | 2186 | $306 / 100 μL |
| Rabbit anti-TOM20 | Cell Signaling Technology | 42406 | $327 / 100 μL |
| Rabbit anti-COXIV | Proteintech | 11242-1-AP-150UL | $189 / 150 μL |
| Mouse anti-TOM20 | Abcam | ab56783 | $283 / 100 μg |
| Mouse anti-COXIV | Cell Signaling Technology | 11967 | $147 / 20 μL |

**Table S3.** List of proteomics raw files used in this report.

| Figure | File Name | Experiment |
|---|---|---|
| 2 | eca10875 | Pan Alpha ISH-POCA |
| 2 | eca10876 | Pan Alpha ISH-POCA |
| 2 | eca10877 | Pan Alpha ISH-POCA |
| 2 | eca10878 | Pan Alpha ISH-POCA |
| 2 | eca10870 | Telomere ISH-POCA |
| 2 | eca10871 | Telomere ISH-POCA |
| 2 | eca10872 | Telomere ISH-POCA |
| 2 | eca10873 | Telomere ISH-POCA |
| 2 | eca11320 | *ITS1* ISH-POCA |
| 2 | eca11321 | *ITS1* ISH-POCA |
| 2 | eca11322 | *ITS1* ISH-POCA |
| 2 | eca11323 | *ITS1* ISH-POCA |
| 2 | eca11310 | *MALAT1* ISH-POCA |
| 2 | eca11311 | *MALAT1* ISH-POCA |
| 2 | eca11312 | *MALAT1* ISH-POCA |
| 2 | eca11313 | *MALAT1* ISH-POCA |
| 3 | 2025-10-09-EB-B-130-NPM1-1 | NPM1 IF-POCA |
| 3 | 2025-10-09-EB-B-130-NPM1-2 | NPM1 IF-POCA |
| 3 | 2025-10-09-EB-B-130-NPM1-3 | NPM1 IF-POCA |
| 3 | 2025-10-09-EB-B-130-SON-1 | SON IF-POCA |
| 3 | 2025-10-09-EB-B-130-SON-2 | SON IF-POCA |
| 3 | 2025-10-09-EB-B-130-SON-3 | SON IF-POCA |
| 3 | 2025-10-09-EB-B-130-Nup98-1 | NUP98 IF-POCA |
| 3 | 2025-10-09-EB-B-130-Nup98-2 | NUP98 IF-POCA |
| 3 | 2025-10-09-EB-B-130-Nup98-3 | NUP98 IF-POCA |
| 3 | 2025-10-09-EB-B-130-NoPrimary-1 | Untargeted (no 1° Ab) IF-POCA |
| 3 | 2025-10-09-EB-B-130-NoPrimary-2 | Untargeted (no 1° Ab) IF-POCA |
| 3 | 2025-10-09-EB-B-130-NoPrimary-3 | Untargeted (no 1° Ab) IF-POCA |
| S8 | 2025-09-13-EB-B-125-COXIV-1 | COXIV IF-POCA |
| S8 | 2025-09-13-EB-B-125-COXIV-2 | COXIV IF-POCA |
| S8 | 2025-09-13-EB-B-125-COXIV-3 | COXIV IF-POCA |
| S8 | 2025-09-13-EB-B-125-Tom20-1 | TOM20 IF-POCA |
| S8 | 2025-09-13-EB-B-125-Tom20-2 | TOM20 IF-POCA |
| S8 | 2025-09-13-EB-B-125-Tom20-3 | TOM20 IF-POCA |
| S8 | 2025-09-13-EB-B-125-NoPrimary-1 | Untargeted (no 1° Ab) IF-POCA |
| S8 | 2025-09-13-EB-B-125-NoPrimary-2 | Untargeted (no 1° Ab) IF-POCA |
| S8 | 2025-09-13-EB-B-125-NoPrimary-3 | Untargeted (no 1° Ab) IF-POCA |
| S8 | 2025-09-25-EB-B-121-CENPA-1-dil | CENPA IF-POCA |
| S8 | 2025-09-25-EB-B-121-CENPA-2-dil | CENPA IF-POCA |
| S8 | 2025-09-25-EB-B-121-CENPA-3-dil | CENPA IF-POCA |
| S8 | 2025-09-25-EB-B-121-Nup98-1 | NUP98 IF-POCA |
| S8 | 2025-09-25-EB-B-121-Nup98-2 | NUP98 IF-POCA |
| S8 | 2025-09-25-EB-B-121-Nup98-3 | NUP98 IF-POCA |

### 3. Supplementary Note

#### General Synthetic Methods

All reactions were carried out in oven-dried glassware using an oven-dried magnetic stir bar under inert atmosphere unless otherwise specified. Reagents and solvents were purchased from reputable suppliers and used without further purification. Analytical thin-layer chromatography (TLC) was carried out using SiliCycle 60Å F254 silica gel (precoated glass-backed sheets, 0.25 mm thickness). Thin-layer chromatography plates were visualized by fluorescence quenching under UV light or by staining with $KMnO_4$, bromocresol green, or ninhydrin. Reaction products were purified by manual silica gel column chromatography using SiliCycle silica gel P60 or by preparative HPLC (Phenomenex Gemini 5 µm NX-C18 110 Å, 150 x 4.6 mm column).

#### General Analytical Methods

$^{1}$H NMR spectra were collected on a 400 MHz or 600 MHz spectrometer in the stated solvents as a reference for internal deuterium lock. $^{13}$C NMR spectra were collected on a Bruker AV400 (101 MHz), AV500 (126 MHz), or NEO600 (151 MHz) spectrometer in the stated solvents as a reference for internal deuterium lock. NMR instruments were provided by the UCLA Molecular Instrumentation Center (MIC) or were located at HHMI Janelia. All chemical shifts are reported as δ in the standard notation of parts per million (ppm) using the peak of residual proton signals of the deuterated solvent as an internal reference. Coupling constant (*J*) units are in Hertz (Hz) to the nearest 0.1 Hz. Splitting patterns are indicated as follows: br, broad; s, singlet; d, doublet; t, triplet; q, quartet; p, pentet; m, multiplet; or combinations thereof. Low-resolution mass spectrometry was performed on an Agilent Technologies InfinityLab LC/MSD single quadrupole LC/MS (ESI source). High-resolution mass spectrometry was performed on a Waters LCT Premier coupled with an ACQUITY LC and autosampler (ESI source) provided by the UCLA MIC or performed by the HHMI Janelia Mass Spectrometry Department.

**6-Carboxy-DBF (4',5'-dibromo-3',6'-dihydroxy-3-oxo-3H-spiro[isobenzofuran-1,9'-xanthene]-6-carboxylic acid).**

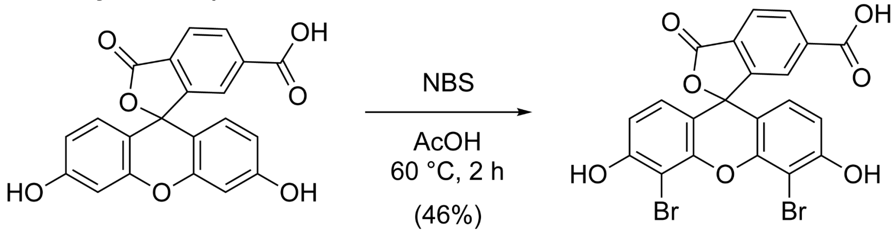


A round-bottom flask was charged with 3',6'-dihydroxy-3-oxo-3H-spiro[isobenzofuran-1,9'-xanthene]-6-carboxylic acid (500 mg, 1.33 mmol, 1 eq.) and acetic acid (40 mL). The yellow suspension was heated to 60 °C while stirring. *N*-Bromosuccinimide (473 mg, 2.66 mmol, 2 eq) was dissolved in acetic acid (20 mL) and added to the reaction mixture dropwise, and the reaction was stirred at 60 °C for 2 h. The crude reaction mixture was then extracted with ethyl acetate (3 x 60 mL). The organic components were combined, washed with brine (150 mL), dried over anhydrous sodium sulfate, and concentrated *in vacuo* to yield a crude red solid. Purification by reversed-phase preparative HPLC (20-100% acetonitrile/$H_2O$ with constant 0.1% FA additive) and lyophilization of product-containing fractions yielded the title compound as a red solid (324 mg, 0.606 mmol, 46%).

**TLC $R_f$ (50% EtOAc/hexanes, 0.1% formic acid):** 0.2

**$^1$H NMR (600 MHz, DMSO-$d_6$):** δ 10.99 (s, 2H), 8.25 (dd, J = 8.0, 1.3 Hz, 1H), 8.13 (d, J = 8.0 Hz, 1H), 7.81 (s, 1H), 6.77 (d, J = 8.8 Hz, 2H), 6.66 (d, J = 8.8 Hz, 2H).

**$^{13}$C NMR (126 MHz, DMSO-$d_6$):** δ 168.1, 166.5, 157.3, 152.6, 149.2, 138.0, 131.6, 129.8, 128.0, 125.8, 125.3, 113.2, 110.8, 98.0, 83.5.

**LRMS *m/z* (ESI$^+$):** calculated for $C_{21}H_{11}Br_2O_7^+$ ([M+H]$^+$): 534.9, found 534.9

**HRMS *m/z* (ESI$^+$):** calculated for $C_{21}H_{11}Br_2O_7^+$ ([M+H]$^+$): 534.8846, found 534.8847

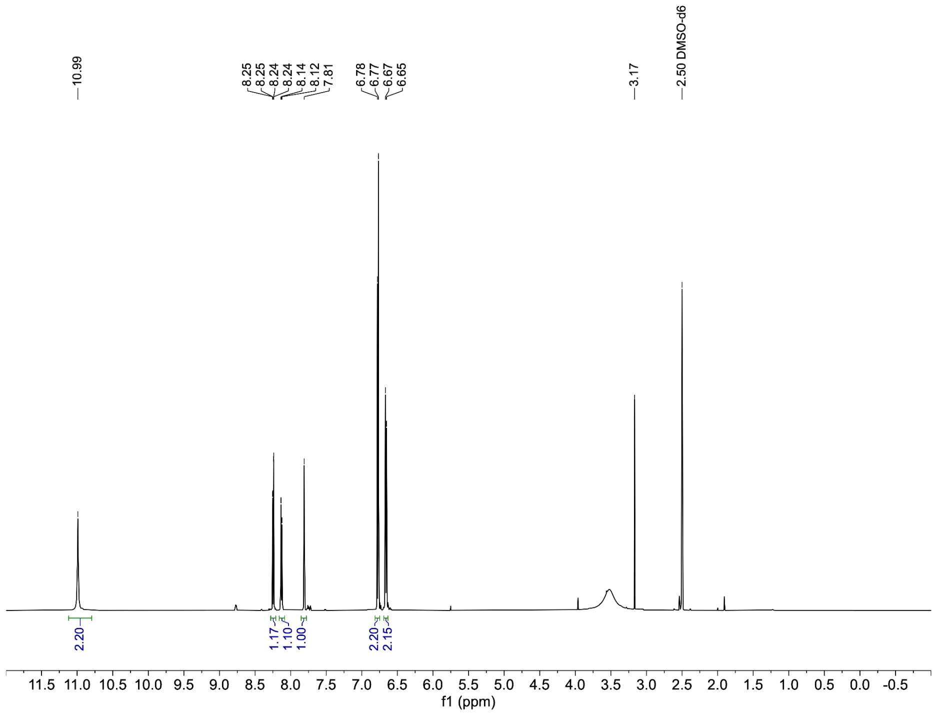
10.99
8.25
8.25
8.24
8.24
8.14
8.12
7.81
6.78
6.77
6.67
6.65
3.17
2.50 DMSO-d6
2.20
1.17
1.10
1.00
2.20
2.15
11.5 11.0 10.5 10.0 9.5 9.0 8.5 8.0 7.5 7.0 6.5 6.0 5.5 5.0 4.5 4.0 3.5 3.0 2.5 2.0 1.5 1.0 0.5 0.0 -0.5
f1 (ppm)

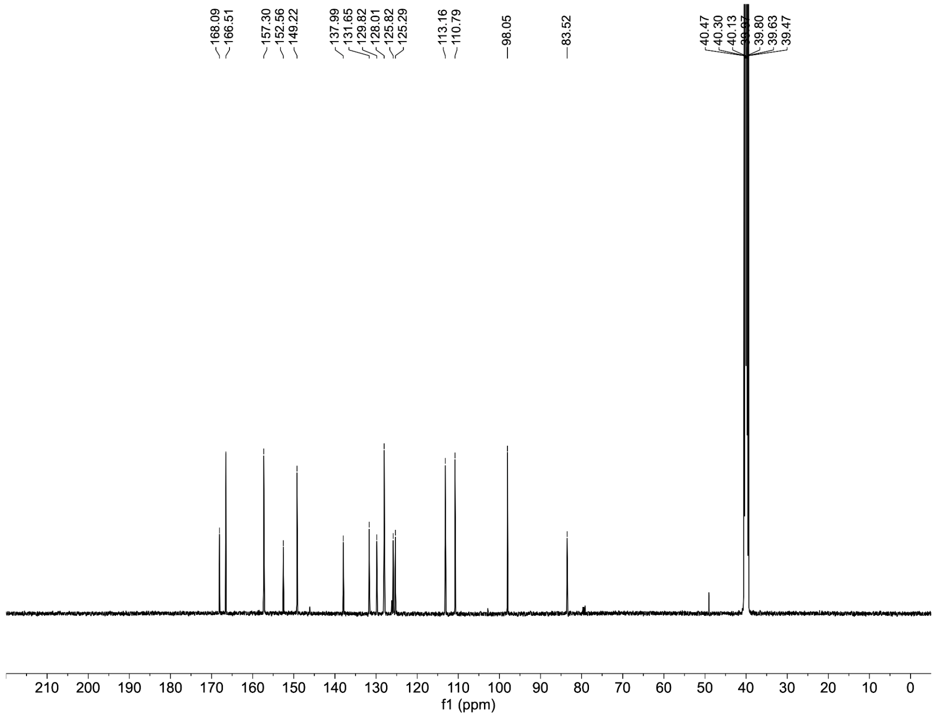
168.09
166.51
157.30
152.56
149.22
137.99
131.65
129.82
128.01
125.82
125.29
113.16
110.79
98.05
83.52
40.47
40.30
40.13
39.80
39.63
39.47
210 200 190 180 170 160 150 140 130 120 110 100 90 80 70 60 50 40 30 20 10 0
f1 (ppm)

**DBF-NHS (2,5-dioxopyrrolidin-1-yl 4',5'-dibromo-3',6'-dihydroxy-3-oxo-3H-spiro[isobenzofuran-1,9'-xanthene]-6-carboxylate).**

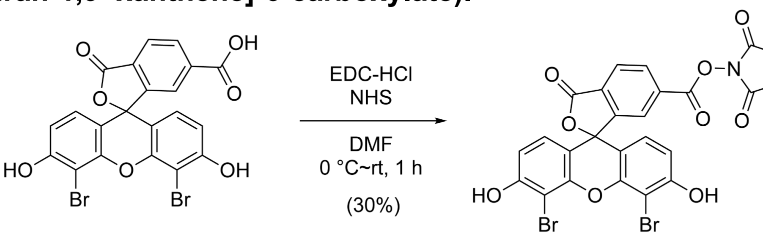


To a microwave vial equipped with a spin vane was added 4',5'-dibromo-3',6'-dihydroxy-3-oxo-3H-spiro[isobenzofuran-1,9'-xanthene]-6-carboxylic acid (50 mg, 0.094 mmol, 1 eq) and DMF (500 μL, anhydrous), and the orange solution was cooled to 0 °C in an ice-water bath. Then, a solution of EDC (20 mg, 0.10 mmol, 1.1 eq) and NHS (12 mg, 0.10 mmol, 1.1 eq) in DMF (500 μL) was added to the reaction mixture. The ice-water bath was removed, the headspace was evacuated and backfilled with argon, and the resulting orange solution was stirred at ambient temperature under inert atmosphere for 1 h. Purification by reversed-phase preparative HPLC (20-100% MeCN/$H_2O$ with constant 0.1% FA additive) and lyophilization of product-containing fractions yielded the title compound as an orange solid (18 mg, 0.029 mmol, 30%).

**TLC $R_f$ (10% MeOH/DCM):** 0.1

**$^1$H NMR (600 MHz, MeOD):** δ 8.41 (dd, J = 8.1, 1.4 Hz, 1H), 8.22 (dd, J = 8.1, 0.8 Hz, 1H), 7.99 (dd, J = 1.4, 0.8 Hz, 1H), 6.77 – 6.65 (m, 4H), 3.35 (s, 1H), 2.9 – 2.8 (m, 4H), 2.85 (s, 4H).

**$^{13}$C NMR (126 MHz, MeOD):** δ 173.5, 170.1, 167.9, 160.8, 149.9, 131.6, 131.2, 127.3, 127.0, 126.3, 126.2, 113.1, 110.9, 98.3, 25.1, 25.0, 24.9

**LRMS *m/z* (ESI$^+$):** calculated for $C_{25}H_{14}Br_2NO_9^+$ ([M+H]$^+$): 631.9, found 631.9

**HRMS *m/z* (ESI$^+$):** calculated for $C_{25}H_{14}Br_2NO_9^+$ ([M+H]$^+$): 631.9010, found 631.9019

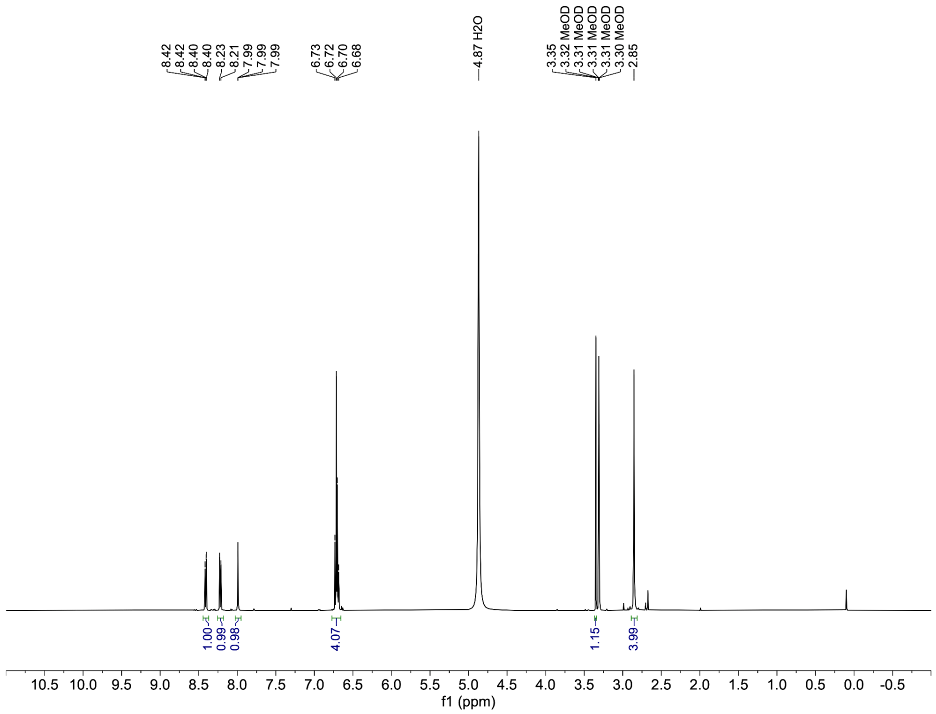
8.42
8.42
8.40
8.40
8.23
8.21
7.99
7.99
7.99
6.73
6.72
6.70
6.68
4.87 H2O
3.35
3.32 MeOD
3.31 MeOD
3.31 MeOD
3.31 MeOD
3.30 MeOD
2.85
1.00
0.99
0.98
4.07
1.15
3.99
f1 (ppm)

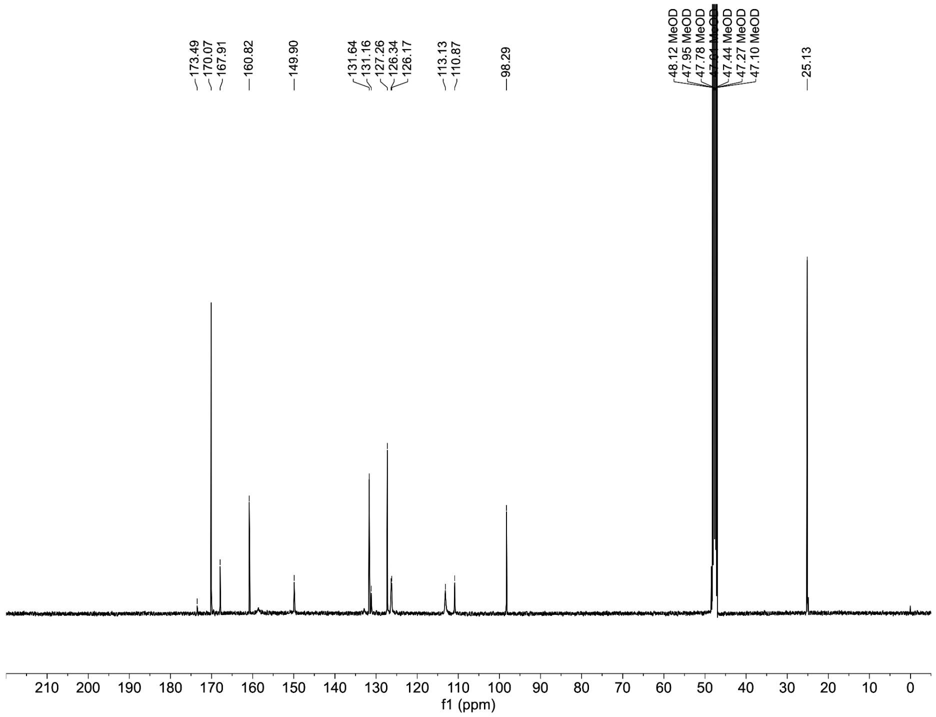
173.49
170.07
167.91
160.82
149.90
131.64
131.16
127.26
126.34
126.17
113.13
110.87
98.29
48.12 MeOD
47.95 MeOD
47.78 MeOD
47.44 MeOD
47.27 MeOD
47.10 MeOD
25.13
f1 (ppm)

**$JF_{570}$-NHS (2-(3-(azetidin-1-ium-1-ylidene)-6-(azetidin-1-yl)-3*H*-thioxanthen-9-yl)-4-(((2,5-dioxopyrrolidin-1-yl)oxy)carbonyl)benzoate).**

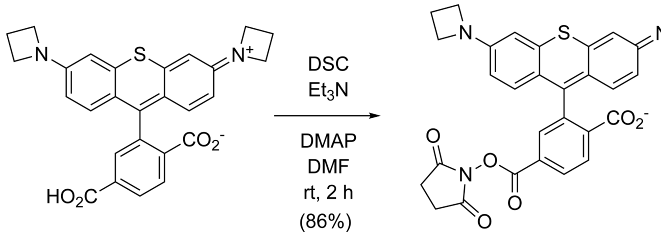


6-Carboxy-$JF_{570}$ [10]; acetate salt; 30 mg, 56.5 μmol, 1 eq.) was combined with *N,N'*-disuccinimidyl carbonate (DSC) (35 mg, 0.14 mmol, 2.4 eq) in DMF (2 mL). After adding triethylamine ($Et_3N$) (47.3 μL, 0.339 mmol, 6 eq) and 4-(Dimethylamino)pyridine (DMAP) (0.7 mg, 6 μmol, 0.1 eq), the reaction was stirred at room temperature for 2 h. The crude reaction mixture was directly purified by reverse phase HPLC (10–75% MeCN/$H_2O$, linear gradient, with constant 0.1% v/v TFA additive) to afford the title compound as a dark purple solid (33 mg, 48 μmol, 86%, trifluoroacetate salt).

**$^1$H NMR (400 MHz, DMSO-*d*$_6$):** δ 8.46 (dd, J = 8.3, 1.7 Hz, 1H), 8.42 (d, J = 8.3 Hz, 1H), 8.05 (d, J = 1.7 Hz, 1H), 7.09 (d, J = 9.4 Hz, 2H), 7.03 (d, J = 2.3 Hz, 2H), 6.64 (dd, J = 9.4, 2.3 Hz, 2H), 4.24 (t, J = 7.6 Hz, 8H), 2.90 (s, 4H), 2.44 (p, J = 7.6 Hz, 4H)

**HRMS *m/z* (ESI$^+$):** calculated for $C_{31}H_{26}N_3O_6S^+$ ($[M+H]^+$): 568.1537, found 568.1538.

**Analytical HPLC:** $t_R$ = 11.2 min, >99% purity (10–75% MeCN/$H_2O$, linear gradient, with constant 0.1% v/v TFA additive; 20 min run; 1 mL/min flow; ESI; positive ion mode; detection at 575 nm)

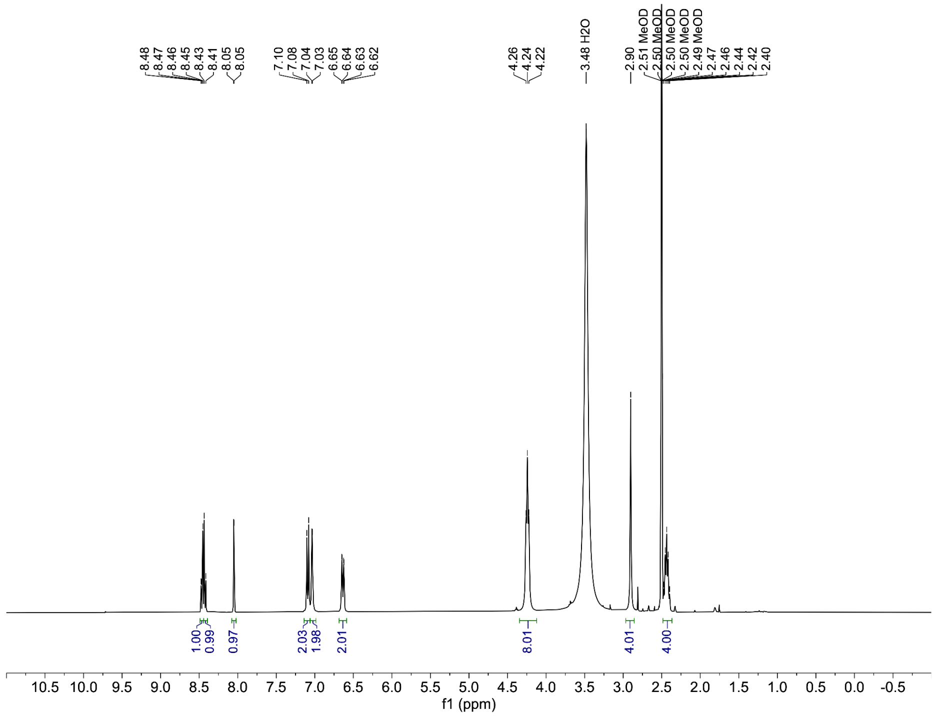
8.48
8.47
8.46
8.45
8.43
8.41
8.05
8.05
7.10
7.08
7.04
7.03
6.65
6.64
6.63
6.62
4.26
4.24
4.22
3.48 H2O
2.90
2.51 MeOD
2.50 MeOD
2.50 MeOD
2.49 MeOD
2.47
2.46
2.44
2.42
2.40
1.00
0.99
0.97
2.03
1.98
2.01
8.01
4.01
4.00
10.5 10.0 9.5 9.0 8.5 8.0 7.5 7.0 6.5 6.0 5.5 5.0 4.5 4.0 3.5 3.0 2.5 2.0 1.5 1.0 0.5 0.0 -0.5
f1 (ppm)

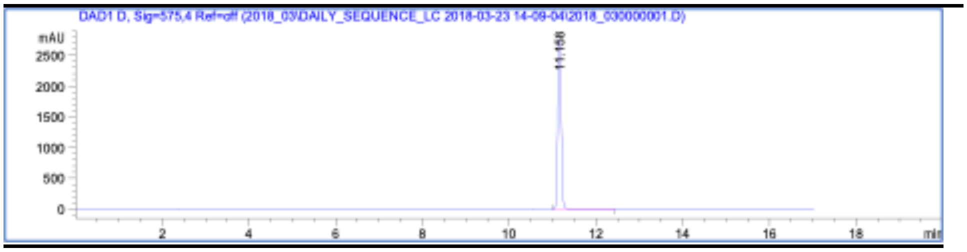
DAD1 D, Sig=575,4 Ref=off (2018_03\DAILY_SEQUENCE_LC 2018-03-23 14-09-04\2018_030000001.D)
mAU
2500
2000
1500
1000
500
0
11.158
2
4
6
8
10
12
14
16
18
min
